# Identification of a new cell cycle variant during multiciliated cell differentiation

**DOI:** 10.1101/2024.05.22.595357

**Authors:** Jacques Serizay, Michella Khoury Damaa, Amélie-Rose Boudjema, Rémi Balagué, Marion Faucourt, Nathalie Delgehyr, Camille Noûs, Laure-Emmanuelle Zaragosi, Pascal Barbry, Nathalie Spassky, Romain Koszul, Alice Meunier

## Abstract

2

A complex and conserved regulatory network drives the cell cycle. Individual components of this network are sometimes used in differentiated cells, i.e. to control organelle destruction in mammalian lens cells or light response in land plants. Some differentiated cells co-opt cell-cycle regulators more largely, to increase their ploidy using a cell cycle variant named endoreplication. Using single-cell RNA-seq profiling and functional assays in differentiating multiciliated cells, we identified a novel type of cell cycle variant that supports cytoplasmic organelle, rather than nuclear content amplification. This variant operates in post-mitotic, centriole-amplifying differentiating multiciliated cells and is characterized by (i) a circular trajectory of the transcriptome, (ii) sequential expression of more than 70% of the genes involved in S, G2 and M-like progression along this trajectory, and (iii) successive waves of cyclins. This cell cycle variant is tailored by the expression of the non-canonical cyclins O and A1 – which replace the transcriptionally silent cyclins E2 and A2 – and by the silencing of the APC/C inhibitor Emi1, two switches also detected in male meiosis, another variant of the canonical cell cycle where centriole and DNA replications are uncoupled. Re-expressing Cyclin E2, cyclin A2 or Emi1 is sufficient to induce partial replication and mitosis, suggesting that change in the regulation of expression of a few cell cycle key players drives a qualitative and quantitative tuning of Cdk activity, allowing the diversion of the cell cycle in the multiciliation variant. We also propose that this new cell cycle variant relies on the existence of a cytoplasmic – or centriolar – Cdk threshold, lower than the S-phase threshold, which affects only the cytoplasmic reorganization.

**One-Sentence Summary:** MCC progenitors undergo a final, tailored iteration of the cell cycle during differentiation, to drive centriole amplification without DNA replication or mitosis.

## 3 Main Text

Centriole duplication and DNA replication processes are regulated by shared factors throughout the cell cycle, and centriole over-duplication is typically associated with genomic instability, accelerating tumor formation and increasing tumor invasiveness (review in (*1*)). Centrioles are also the essential component of basal bodies, which act as anchors from which cilia nucleate. Multiciliated cells (MCCs) harbor hundreds of cilia to transport fluids along organ lumens and promote essential respiratory, reproductive, and brain functions. To sustain basal body production required for the formation of hundreds of cilia, post-mitotic MCC progenitors need to uncouple centriole biogenesis from DNA replication to massively amplify centrioles. In the mouse brain, it occurs following three stereotypical phases: the centriole amplification phase in which centrioles are massively amplifying around “deuterosome” organelles, the centriole growth phase in which centrioles grow, and finally the centriole disengagement phase in which the newly formed centrioles will migrate and dock to the apical plasma membrane to act as molecular anchors for the future cilia (*2*).

Despite major differences with centriole duplication such as the massive production of centrioles and the intervention of MCC specific organelles, we and others have previously shown that the activity of individual cell cycle key players such as MYB, CDK2, CDK1, PLK1 and the APC/C was essential for accurate control of centriole amplification in MCCs (*3–8*). Pharmaceutical modulation of the activity of these cell cycle factors induces major defects in numbers of generated centrioles and in the dynamics of their formation, as well as in motile ciliation in terminally differentiated MCCs. They can also induce mitosis-like features and abnormal DNA replication (*3*, *5*). Here we use single-cell RNA-seq experiments to investigate the co-option of cell cycle factors, coupled with functional assays to study the role of cyclins during MCC differentiation. We reveal that MCC differentiation is controlled by a genuine cell cycle variant where CCNO and CCNA1, non-canonical cyclins associated with the expression of genes involved in centriole amplification and motile ciliation, respectively, replace CCNE2 and CCNA2 cyclins canonically involved in DNA synthesis and mitotic division. We further show that this cell cycle variant is tailored by a limited expression of *Emi1* APC/C inhibitor. Rescuing the expression of canonical cyclins and/or EMI1 APC/C inhibitor is sufficient to reroute some differentiating nuclei into partial DNA replication and/or nuclear division. In a companion study, we further show that *Ccno*, mutated in patients with a severe congenital ciliopathy, is a key regulator of MCC differentiation regulating the entry into this cell cycle variant (**Khoury Damaa et al., 2024, ref. 44**). These results show that MCC differentiation is a final and tailored iteration of the cell cycle allowing centriole amplification without DNA replication or mitosis.

## Results

### Post-mitotic differentiating MCC massively express cell-cycle factors during differentiation

We sought to investigate the extent to which cell cycle factors are co-opted in MCC differentiation. We harvested MCC progenitor cells from lateral ventricle walls of P1 mouse brains and cultured cycling progenitors to confluence before inducing their differentiation into MCCs (see Methods). We then profiled the single-cell transcriptomes of 16,401 cells comprising radial glial progenitors during and after they exit proliferation phase, differentiating MCC progenitors (also known as deuterosomal cells) and terminally differentiated MCCs (**Fig. 1A**, **Fig. S1A**). After removing contaminating cells (e.g. oligodendrocytes, neuroblasts, fibroblasts) and correcting for batch effects (**Fig. S1B**), we annotated four well-defined cell populations subdivided into eight clusters: cycling progenitors (expressing *Mki67*), post-mitotic progenitors (2 clusters marked by *Id1* and *Id3* expression but no *Mki67*), deuterosomal cells (3 clusters: early (marked by *Deup1* and *Ccno*), mid (marked by max *Deup1* expression) and late (marked by *Deup1* and *Ube2c*) deuterosomal cells) and multiciliated cells (2 clusters: early (marked by *Ccna1* expression but no *Ube2c*) and terminal (marked by *Tmem212*) MCCs) (**Fig. 1B**, **Fig. S1C**, **Table S1**, see Methods). The remaining 2,534 unannotated cells showed no clear overexpression of marker genes and were characterized by low clustering stability (**Fig. S1, D-E**).

**Fig. 1:**
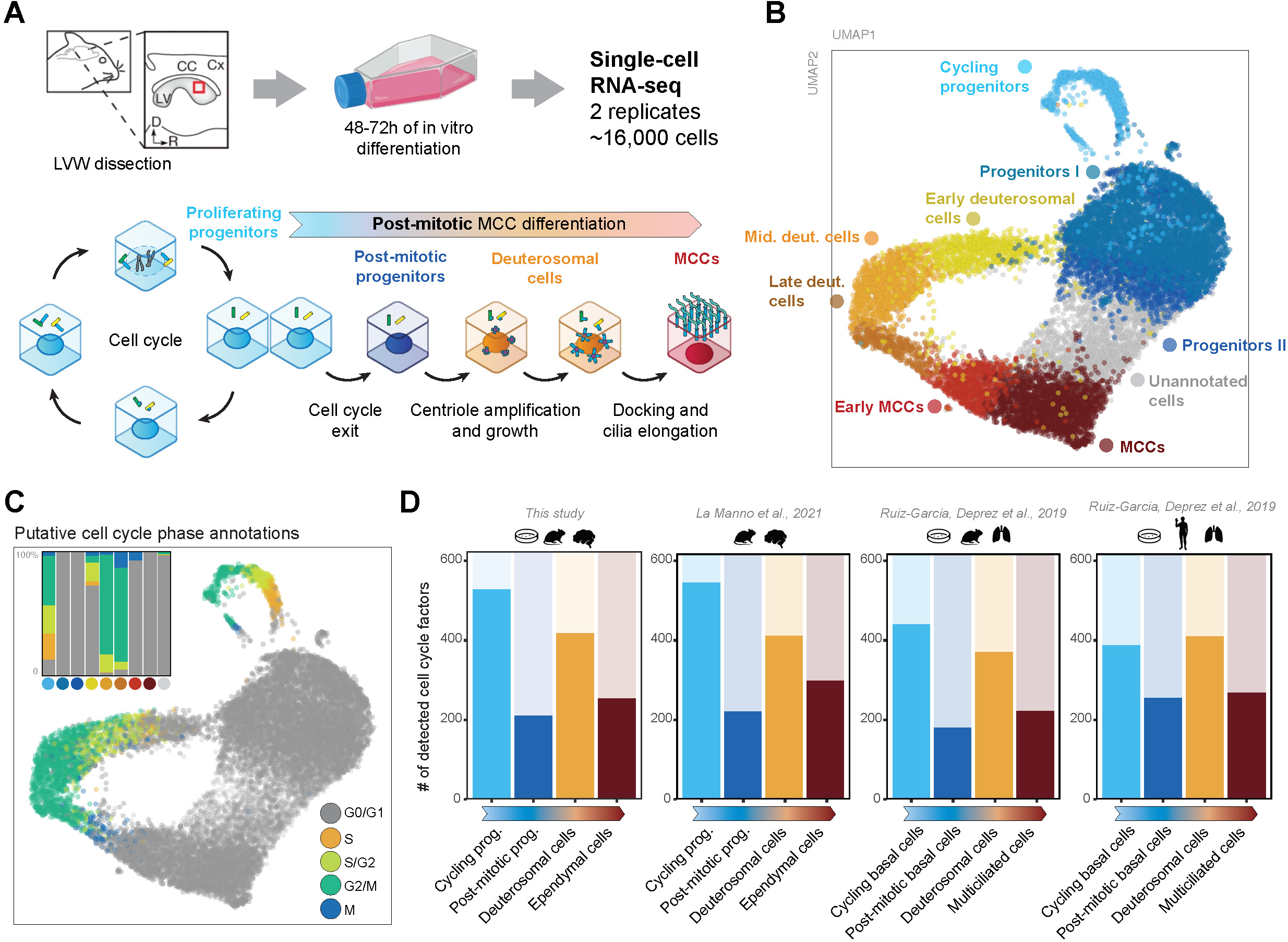
Deuterosomal cells massively co-opt cell-cycle factors during post-mitotic differentiation. **(A)** Single-cell RNA-sequencing profiling of in vitro mouse radial glial cells differentiating into multiciliated cells. **(B)** UMAP projection of the single-cell RNA-seq dataset. Colors represent annotated cell populations. Post-mitotic progenitor cells, deuterosomal cells and MCCs have been further divided in smaller cell clusters. **(C)** Putative cell cycle phase annotations inferred using SingleR using neural stem cell reference (*9*). The inset represents the proportion of each putative cell cycle phase in each cluster. **(D)** Number of cell cycle–related factors detected in cycling progenitors, post-mitotic progenitors, deuterosomal cells and multiciliated cells. Single-cell RNAseq experiments performed in vitro or in vivo from mouse and human brain or tracheal samples (*11*, *12*) all reveal expression of a large number of cell cycle–related factors in deuterosomal cells. Cell cycle factors were retrieved from (*10*).

Using cell cycle phase transcriptional signatures from a neural stem cell single-cell RNA-seq reference (*9*), we annotated putative cell cycle phases for each cell. Interestingly, we found most deuterosomal cells (70%) were annotated in either S, G2 or M phases, especially in the later deuterosomal stage (**Fig. 1C**, **Fig. S1, F-G**). We leveraged a list of manually curated cell cycle-related genes (n = 623 mouse genes) (*10*) to investigate their co-option during differentiation. Whereas 85% (527/623) of the cell cycle genes are expressed in cycling progenitors, only 34% (210/623) remained expressed in post-mitotic progenitors (**Fig. 1D**). These numbers are comparable to those observed between proliferating oligodendrocyte progenitors (OPCs) and post-mitotic oligodendrocytes (**Fig. S1H**). However, as much as 67% (417/623) of the cell cycle genes were also expressed in deuterosomal cells, while only 41% (253/623) remained expressed in post-mitotic MCC (**Fig. 1D**). This highlights that hundreds of cell cycle genes are reactivated during MCC differentiation, specifically at the deuterosomal stage during which centrioles are amplifying. We observed comparable distribution of cell cycle phase transcriptional signatures and a reactivation of cell cycle factors in deuterosomal cells profiled from in vivo mouse embryonic brain development (*11*) as well as from respiratory cells in mouse and human (*12*) (**Fig. 1D, Fig. S2**), showing that this is not specific to brain or cultured mouse MCCs and is conserved in Human. We validated this observation using a cluster-independent differentiation trajectory analysis (**Fig. S3**). Altogether, these results reveal an unexpected and massive co-option of cell cycle genes during MCC differentiation, across species and tissues, resulting in transcriptional signatures of deuterosomal cells similar to that of cycling cells in different cell cycle phases.

### Factors from most cell cycle-related molecular functions are expressed in deuterosomal cells

We next aimed at precisely determining in which cell cycle subprocesses the genes that are reexpressed in deuterosomal cells were specifically involved. Using the cell cycle factor classification from Giotti et al. 2019, we observed that genes expressed in deuterosomal cells are significantly enriched amongst 15 of 17 cell cycle-related sets of genes (**Fig. 2, A-B**). As expected, nearly all genes related to centrosome regulation (94%, 32/34) are strongly detected in deuterosomal cells (**Fig. S4A**). Interestingly, factors involved in all the other cell cycle-related processes are also widely co-opted (e.g. all of the 19 genes involved in cytokinesis, 15 out of 19 genes involved in nuclear envelope regulation, etc.) (**Fig. S4A**), suggesting that beyond centriole biogenesis, all cellular compartments may be affected by cell cycle-like reorganization during MCC differentiation. Notably, 59% of the genes directly involved in DNA replication (30/51) are unexpectedly robustly re-expressed in deuterosomal cells (e.g. catalytic and regulatory subunits of the DNA replication polymerase *Pola1*, *Pole3*) (**Fig. S4A**). Similarly, a significant number of cell cycle factors involved in DNA condensation, chromosome partition, kinetochore formation (8/11, 19/26, 22/25 respectively; **Fig. S4A**) like condensin subunits (e.g. *Smc2*, *Smc4*, *Ncap* proteins, etc.), topoisomerase 2A (*Top2a*) or *CenpA* are unexpectedly reexpressed in deuterosomal cells. Such recruitment of nuclear factors that apparently have no reason to be expressed in MCC suggests the existence of cell cycle-like events in the nucleus during differentiation and/or may indicate unknown functions of these factors in the cytoplasm.

**Fig. 2:**
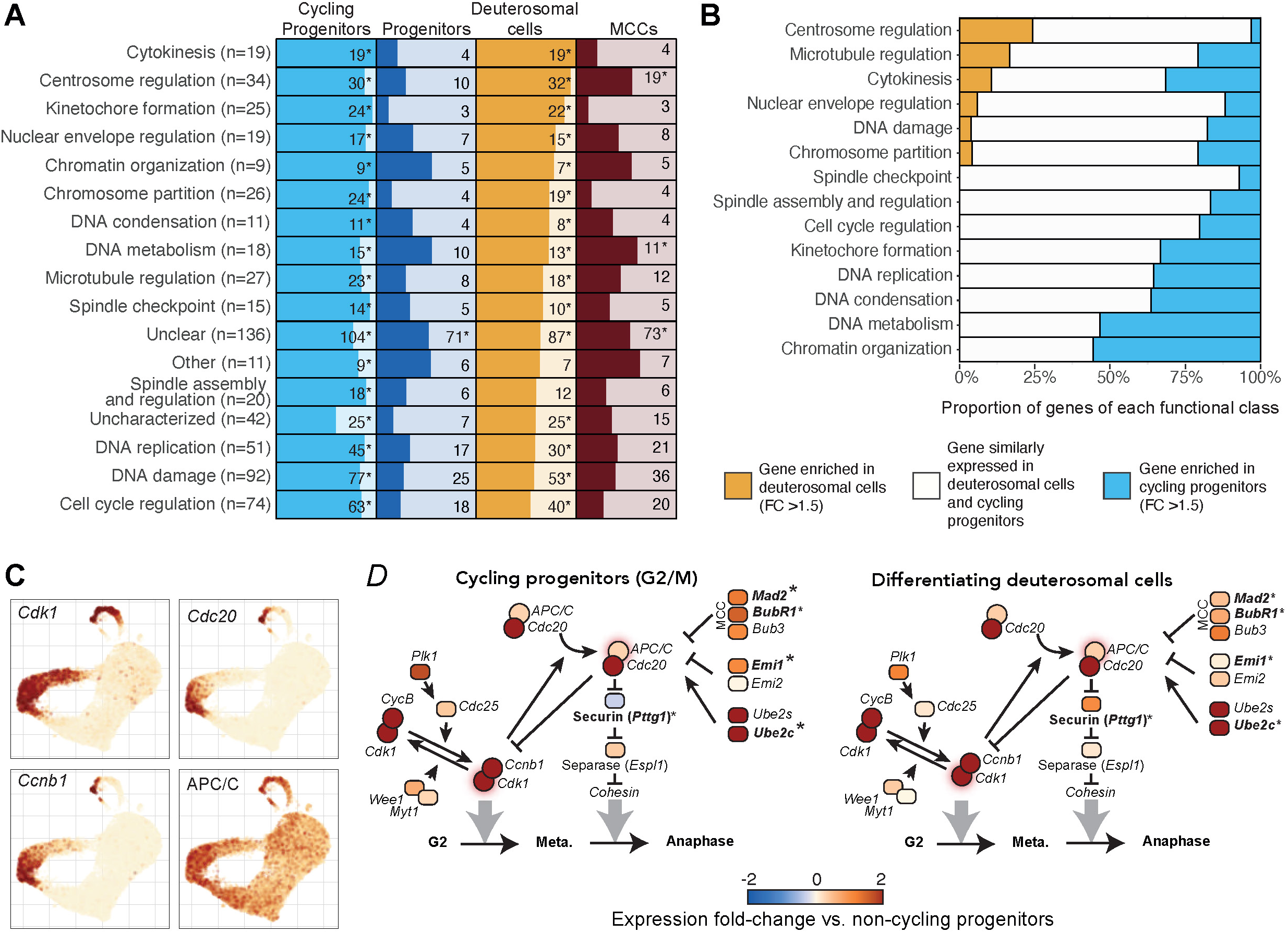
Factors from all cell cycle subprocesses are re-expressed in deuterosomal cells. **(A)** Table of cell cycle factors expressed in each cell population. Cell cycle factors were manually classified by their main function in the cell cycle (*10*). Stars denote cell cycle functions statistically enriched (> 4-fold) for genes expressed in cycling progenitors, post-mitotic progenitors, deuterosomal cells or in MCCs. The total number of factors in each functional class is indicated in parentheses. **(B)** Proportion of cell cycle factors preferentially enriched in deuterosomal cells versus cycling progenitors (orange), or in cycling progenitors versus deuterosomal cells (in blue), or not differentially expressed (in white). Genes which are not detected in any of the two cell populations are not included. **(C)** UMAP projection of the scRNAseq dataset, each cell colored by the level of expression of *Cdk1*, Cyclin B (*Ccnb1*), *Cdc20* or the average expression of all APC/C subunits. **(D)** Schematic of the main components of the mitotic oscillator in G2/M cycling progenitor cells (top) or in deuterosomal cells (bottom). The colormap indicates gene expression fold-change versus non-cycling progenitor cells. Labels in bold indicate factors over-expressed between deuterosomal cells and G2/M cycling progenitors (fold-change > 1.5, p-value < 0.01).

Because we previously showed that an attenuated activity of the mitotic oscillator (CDK1 and APC/C) itself was required during MCC differentiation (*3*), we compared gene expression of the mitotic oscillator components and their direct regulators in cycling progenitors or deuterosomal cells. We found that in addition to *Cdk1*, *Ccnb1*, APC/C subunits and *Cdc20*, nearly all mitotic oscillator components are reexpressed in deuterosomal cells to levels significantly comparable to those in cycling progenitors (**Fig. 2, C-D**) showing that the whole machinery is co-opted and refuting the hypothesis that mitotic clock attenuation in MCC is driven by a global decreased expression of the mitotic oscillator components. However, several APC/C inhibitors are reexpressed in differentiating MCCs at lower levels than in cycling progenitors, and the APC/C inhibitor *Emi1/Fbxo5* is totally silenced (see **Fig. 2, C-D, Fig. S4B**) and is replaced by its homolog *Emi2/Fbxo43*, albeit expressed at very low level (**Fig. S4C**), consistent with a role of *Emi2* in MCC differentiation previously documented in Xenopus (*5*). This indicates that an overactive APC/C, rather than a decreased expression of the mitotic oscillator main components, might attenuate the mitotic clock in MCC.

Altogether, these results show that a large proportion of genes from all the cell division processes are reexpressed in differentiating mouse MCC and therefore suggest that, in addition to centriole biogenesis, cell cycle-like reorganization of the cytoplasm, the nuclear membrane, and even the chromatin is occurring during MCC differentiation. Further studies are needed to highlight the other aspects of the cell cycle that are co-opted to control MCC differentiation.

### Deuterosomal cells progress through differentiation by following a transcriptional circular trajectory similar to the cell cycle

The expression of the cell cycle regulatory circuitry in deuterosomal cells suggests that they can progress through the different intermediate stages of differentiation by using regulatory mechanisms thought to be restricted to cycling cells progressing through different cell cycle phases. Supporting this, when processed together, deuterosomal cells cluster with cycling progenitors according to their putative cell cycle phase (**Fig. S5A-D**).

Several reports recently showed that the stereotypical clocklike progression of cells through the cell cycle can be identified from single-cell RNA-seq data (*13–15*). Schwabe et al. and Zinovyev et al. both revealed that immortalized cycling cells embedded in a linear PCA space form a circular path along which they advance as a cycle (*14*, *15*). PCA integration of our primary cycling progenitors (n = 759 cells) highlighted that these cycling primary cells also form a planar circular path in linear 2D space (**Fig. 3A**). We computed a radial progression for each cell, which we used to reveal that cell cycle phases are sequentially distributed along this circular path (**Fig. 3B**), confirming previously published results in immortalized cycling cells. We then performed the same analysis on deuterosomal cells and observed that they recapitulate a similar circular path, successively traversing early, mid and late deuterosomal transcriptional stages (**Fig. 3C**, **Fig. S5E**). Importantly, we found that putative cell cycle-like phase annotations for deuterosomal cells are also sequentially distributed along the circular trajectory (**Fig. 3D**), as shown for cycling progenitor cells. Deuterosomal cells radially progress through G0/G1, S, G2, and M-like phases, albeit with a relative enrichment of S/G2 and G2/M-like cells over pure S-like cells, when compared to the cycling progenitors. This lower representation of S-like cells is consistent with the lower number of DNA replication factors being re-expressed in deuterosomal cells (**Fig. 2A, 2B, S4A**).

**Fig. 3:**
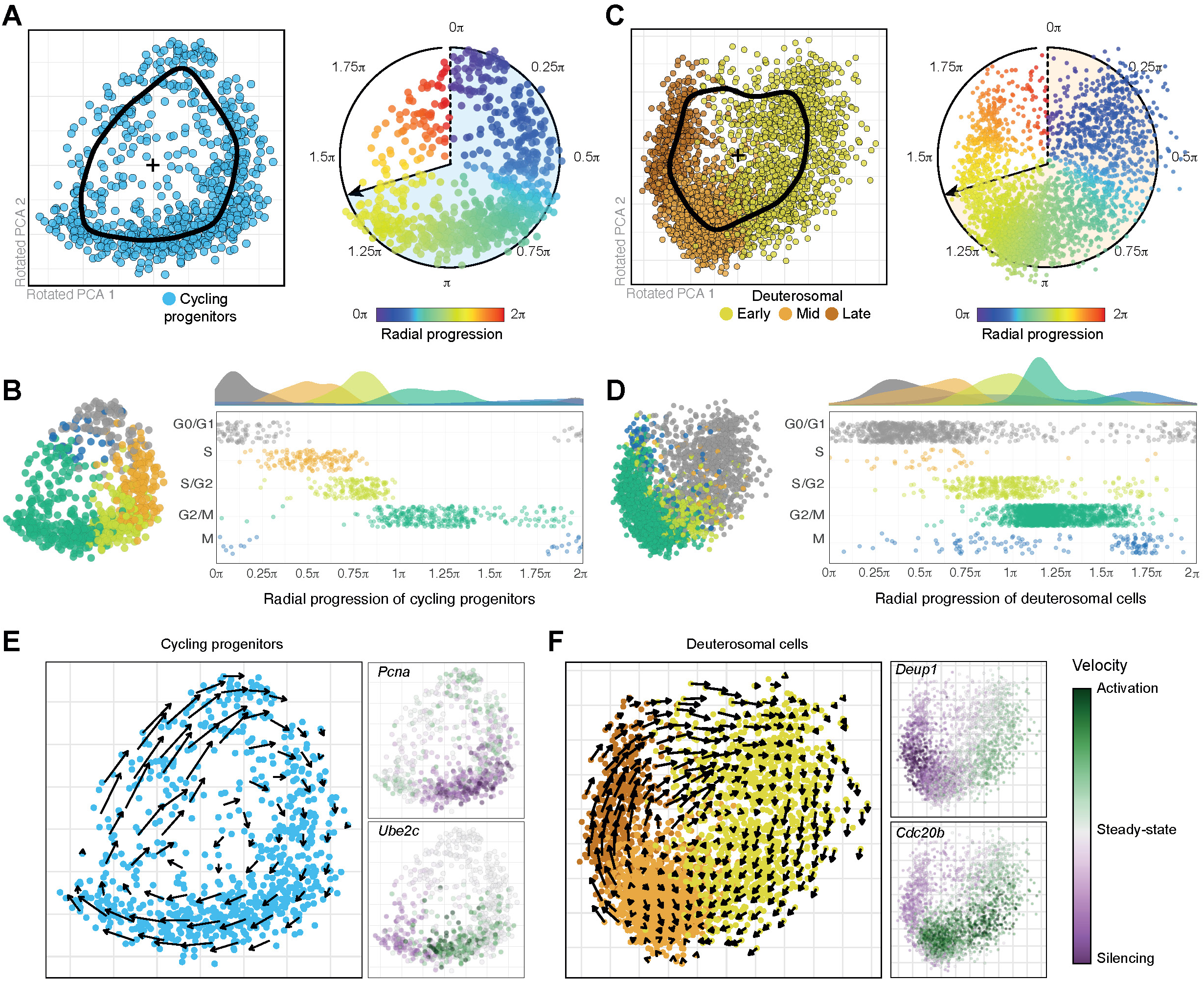
Deuterosomal cells progress through differentiation following a cell cycle-like circular trajectory. **(A)** PCA embedding (PC1 and PC2) of cycling progenitor cells. The black curve denotes the average position of cells around the center point of the PCA space. Right: representation of the cycling progenitors with a color range indicating the radial progression of each cell. **(B)** Radial distribution of cycling progenitors with a color scale indicating the putative cell cycle phase annotated using SingleR and a neural stem cell reference (*9*). **(C)** PCA embedding (PC1 and PC2) of deuterosomal cells. The black curve denotes the average position of cells around the center point of the PCA space. Right: representation of the deuterosomal cells with a color range indicating the radial progression of each cell. **(D)** Radial distribution of deuterosomal cells with a color scale indicating the putative cell cycle phase annotated using SingleR and a neural stem cell reference (*9*). **(E)** RNA velocity analysis of cycling progenitors and deuterosomal cells. RNA velocity relies on nascent transcript abundance to predict where each cell would be positioned in the future. Left: RNA velocity vector field of cycling progenitor cells. Right: RNA velocity score for *Pcna* and *Ube2c*, two cell cycle-related genes. Green (purple) indicates an increasing (decreasing) production of nascent RNA, while white indicates a steady-state production of nascent RNA. **(F)** Left: RNA velocity vector field of deuterosomal cells. Right: RNA velocity score for *Deup1* and *Cdc20b*, two genes involved in the regulation of MCC differentiation.

We then computed RNA velocity in each subset of cells (*16*). In both cycling progenitors and deuterosomal cells, the RNA velocity vector field reveals that cells move forward (i.e. clockwise) along their circular path (**Fig. 3, E-F**). RNA velocity scores of genes transiently transcribed during the cell cycle (e.g. *Cdt1* or *Ube2c*) or MCC differentiation (*Deup1* and *Cdc20b*) confirmed that transcription of these genes sequentially occurs along the respective cycling and deuterosomal circular paths.

In agreement with what has been previously described in immortalized cell lines (*14*) and with our own measurements in cycling progenitors, we also observed a gradual increase in (i) the number of expressed genes and (ii) the total transcript content in deuterosomal cells during the S and G2-like phases, up to a tipping point after which transcription is reduced, as seen in cycling cells after mitosis entry. Also, similarly to what has been previously described in immortalized cycling cell lines (*14*) and to what we observe in primary cycling progenitors, we found two regimes of gene transcription onsets in deuterosomal cells: first a relatively fast rate for 96% of the genes variably expressed in deuterosomal cells (2,106/2,183), followed by a stark decrease (∼4.5-fold decrease) shortly before the cells reach their maximum transcriptional output (**Fig. S6B**).

Altogether, these results show that brain MCC differentiating cells share all the features of transcriptional regulation previously characterized in cycling cells. They follow a multiciliation trajectory characterized by successive cell cycle-like phases and reuse the optimized principles of cell cycle gene regulation. We observed a similar distribution of cell cycle-like transcriptional signatures along a circular trajectory of deuterosomal cells during in vivo mouse embryonic brain development (*11*) (**Fig. S7A-C**). We therefore conclude that the differentiation lineage of MCC consists of a single iteration of a variation of the cell cycle.

### The MCC cell cycle variant is characterized by a cascade expression of cyclins

Cyclins are the key factors controlling the progression of cycling cells through the cell cycle. When a cyclin accumulates to a sufficient threshold, it acts as a molecular switch that triggers the enzymatic activity of specific cyclin-dependent kinases. It has been established that (i) D-type cyclins are involved in progression through the G1 phase, (ii) E-type cyclins contribute to the transition from G1 to S phase, (iii) Cyclin A2 orchestrates the interphase progression as well as the G2/M transition, (iv) and Cyclin B1 controls progression from entry to exit of mitosis (*17*). Successive waves of cyclin expression provide a tentative model to organize the MCC cell cycle variant.

Leveraging the cyclic progression of proliferating progenitor cells, we modeled the temporal activation of canonical cyclins in cycling cells, which recapitulated the well-known expected pattern of successive waves of cyclin transcription, with D2, E2, A2 and B1 cyclins broadly delineating the different putative cell cycle phases (**Fig. 4, A-B**). This result confirms that the radial distribution of dividing cells along a single-cell RNA-seq circular trajectory can be used to estimate cell cycle progression in proliferating progenitors (**Fig. 4B**). Then, using the same approach in deuterosomal cells, we found that Cyclin D2 (*Ccnd2*) and B1 (*Ccnb1*) were transiently expressed in the MCC cell cycle variant (**Fig. 4, C-D**). The temporal expression profiles of these two cyclins were highly correlated with those from cycling progenitors (r=0.98 for *Ccnd2* and r=0.97 for *Ccnb1*), albeit an expression generally lower for *Ccnd2* (**Fig. 4D**). In contrast, the other canonical cyclins (i.e. A2 and E-type cyclins) remained mostly untranscribed throughout this cell cycle variant. Instead, Cyclin O (*Ccno*) and A1 (*Ccna1*), previously shown to be involved in MCC differentiation (*4*, *18*, *19*), are as highly expressed as *Ccnd2* and *Ccnb1* in the MCC cell cycle variant (**Fig. 4, C-D**, **Fig. S8A**).

**Fig. 4:**
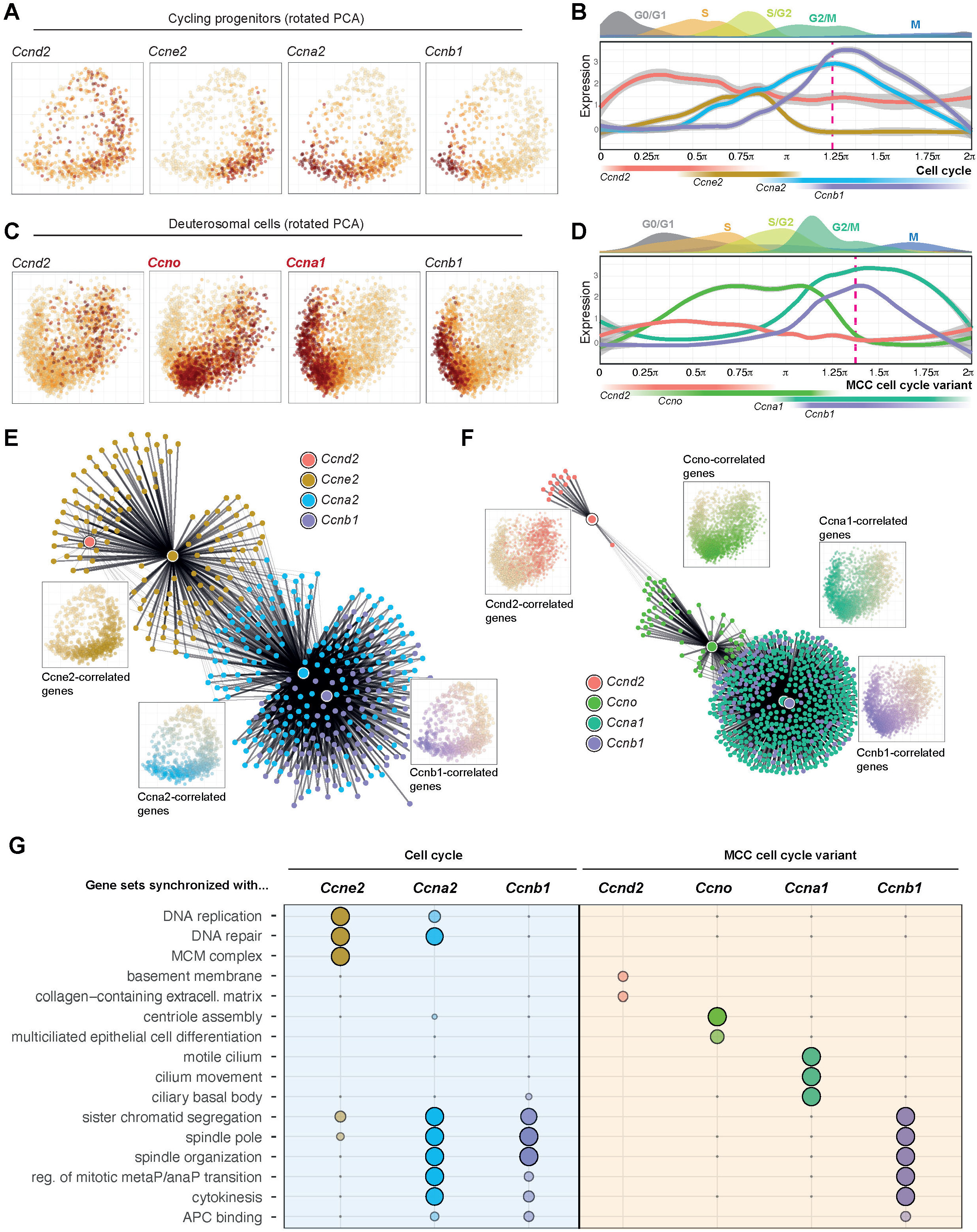
Deuterosomal cells activate cascades of canonical and non-canonical cyclins with coordinated gene expression. **(A)** Expression of Cyclin D2 (*Ccnd2*), Cyclin E2 (*Ccne2*), Cyclin A2 (*Ccna2*) and Cyclin B1 (*Ccnb1*) canonical cyclins in cycling progenitor cells in PCA embedding. **(B)** Expression of these cyclins in cycling progenitors distributed according to their radial progression. This recapitulates the well-known pattern of successive waves of cyclin expression during the cell cycle. **(C)** Expression of Cyclin D2 (*Ccnd2*), Cyclin O (*Ccno*), Cyclin A1 (*Ccna1*) and Cyclin B1 (*Ccnb1*) cyclins in deuterosomal cells in PCA embedding. **(D)** Expression of these cyclins in deuterosomal cells distributed according to their radial progression. Note the successive waves of *Ccnd2*, *Ccno*, *Ccna1* and *Ccnb1* expression, resembling the waves of canonical cyclin expression in cycling progenitors. **(E)** Network of genes (small nodes) whose temporal expression correlates with that of canonical cyclins (larger nodes) in cycling progenitor cells. The colors indicate which cyclin each gene is mostly correlated with. Edge thickness is proportional to the correlation score between gene expression and cyclin expression. Only correlations greater than 0.3 (Pearson ρ score) are shown. Insets show the average expression of all the genes correlated with each cyclin during the canonical cell cycle. **(F)** Network of genes (small nodes) whose temporal expression correlates with that of cyclins (larger nodes) in deuterosomal cells. The colors indicate which cyclin each gene is mostly correlated with. Edge thickness is proportional to the correlation score between gene expression and cyclin expression. Only correlations greater than 0.3 (Pearson ρ score) are shown. Insets show the average expression of all the genes correlated with each cyclin during the MCC cell cycle variant. **(G)** Biological functions associated with sets of genes temporally correlated with each cyclin, in cycling progenitors (left) or in deuterosomal cells (right). Genes with a ρ correlation score greater than 0.3 for several cyclins were associated with the cyclin for which they had the greatest correlation.

Importantly, we uncovered that their expression patterns were temporally delimited, defining two different periods for which each cyclin is predominantly expressed. *Ccno* is prominently expressed from the S and up to the G2/M-like phase of the MCC cell cycle variant, while *Ccna1* is mostly expressed during the G2/M and M-like phases (**Fig. 4D**). In addition, the temporal expression patterns of *Ccno* and *Ccna1* in deuterosomal cells appear correlated with those of *Ccne2* and *Ccna2* in cycling progenitors, respectively (r=0.83 and 0.62). Importantly, we observed a similar temporality of successive cyclin expression along MCC differentiation, both during in vivo mouse embryonic brain development (**Fig. S8B**) (*11*) and in the human epithelial airway (**Fig. S8C**) (*20*). Altogether, these results show that MCC differentiation is marked by successive, partially overlapping, waves of cyclin transcription, analogous to those described in the cell cycle, with sequential D2, O, A1 and B1 cyclin phases, further supporting the existence of a genuine cell cycle variant during MCC differentiation.

Waves of cyclin expression segment the cell cycle into successive phases. In each phase, functional sets of genes are predominantly expressed to carry out specific biological functions in each cell cycle phase (*21*, *22*). Indeed, we found hundreds of genes whose temporal expression was positively correlated with that of canonical cyclins within the cell cycle (n=130 genes with highest correlation for Cyclin E2, n=187 for Cyclin A2, n=110 for Cyclin B1) ( **Fig. 4E**). The functions of these gene sets are respectively enriched for replication processes (MCM complex, nuclear DNA replication), mitosis regulation (e.g. centrosome regulation, spindle checkpoint) and late mitosis events (e.g. APC/C activity), confirming that cyclin-correlated patterns of temporal expression allow defining groups of genes that are functionally related to the cyclin (**Fig. 4G**). The same analysis performed on deuterosomal cells pointed at hundreds of genes with expression positively correlated with that of Cyclin O (n=60), Cyclin A1 (n=499) or Cyclin B1 (n=115). Genes mostly correlated with Cyclin B1 were involved in mitosis events, which some components have been involved in centriole number, growth and disengagement (*2*) (**Fig. 4G**). Genes mostly correlated with Cyclin O appear enriched for key multiciliated cell differentiation and centriole biogenesis regulators. Those mostly correlated with Cyclin A1 are enriched for cilium biogenesis and motility, which contradicts recent data suggesting an early involvement of this CCNA1 in centriole amplification (*4*). Interestingly, we found the same consecutive expression of *Ccno* and *Ccna1* and the same switch between *Emi1* and *Emi2* expression during male meiosis (*23*) (**Fig. S8, D-E**), another important cell cycle variant during which centriole biogenesis occurs -in the absence of DNA replication- and is followed by motile flagella growth.

Altogether, these results support the existence of a MCC-specific cell cycle variant, supported by successives waves of cyclin expression, in which the cell cycle is deviated from its normal trajectory by the replacement of some canonical cyclins by non canonical ones. In this variant conserved across tissues and mammals, the onset is marked by the expression of *Ccno* instead of *Ccne2*, consistent with an increase in cytoplasmic centriole content instead of nuclear ploidy. Later, and with the same temporality as in the cell cycle, *Ccnb1* is expressed, consistent with the role of the mitotic oscillator in centriole growth and disengagement (*3*). During a comparable period, *Ccna1* is expressed instead of the second mitotic cyclin, *Ccna2*, and its expression is correlated with genes involved in cilia motility.

### Expression of canonical cyclins E2, A2 and/or APC/C inhibitor EMI1 are sufficient to trigger cell cycle nuclear events

We next hypothesized that if MCC differentiation is a true cell cycle variant, re-expression of the missing canonical elements would restore the nuclear events skipped during differentiation. In particular, *Emi1* – an APC/C inhibitor that regulates the progression of the canonical cell cycle by regulating DNA replication and its coupling to mitosis (*24–26*) is drastically silenced during MCC differentiation and partially replaced by its homolog *Emi2*, lowly expressed in deuterosomal cells (**Fig. 2D, S4B, S4C**). We should be able to restore DNA replication and/or mitotic events specifically in differentiating cells by re-expressing missing EMI1. Supporting this hypothesis, a recent study in Xenopus reported EdU incorporation and mitosis figures in differentiated skin multiciliated cells upon APC/C inhibition by overexpression of *Emi2* (*5*). We infected differentiating cells to express *Emi1* and monitored DNA replication – by EdU incorporation – and mitosis entry – by immunostaining phosphorylation of Histone 3 Serine 10 (H3S10p) (see Methods) (**Fig. 5A, 5B**). We observed a partial replication in a subset of differentiating progenitors at 96hpi, which we failed to detect upon infection with GFP or with *Emi1* at 24hpi. We also detected a significant proportion of differentiating MCCs in the mitosis entry figures upon *Emi1* infection, both at 24hpi and at 96hpi.

**Fig. 5:**
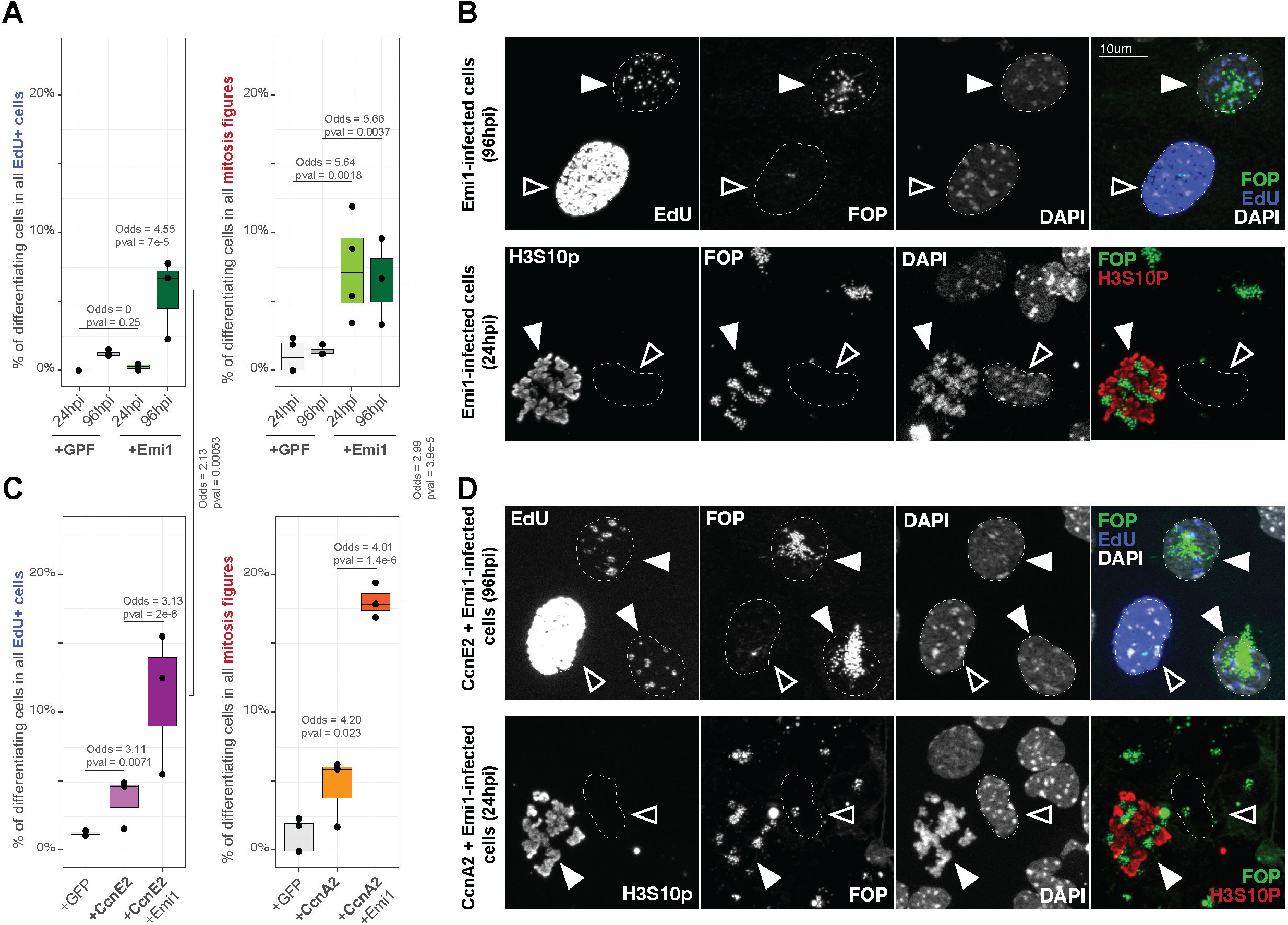
Reactivating APC/C inhibitor *Emi1* or the canonical cyclins E2 or A2 rescues cell cycle nuclear events. **(A)** Proportion of differentiating deuterosomal cells amongst all the cells marked by EdU (left) or mitosis figures (right), after *GFP* or *Emi1* over-expression for 24h and 96h. Differentiating deuterosomal cells are identified according to FOP staining. **(B)** Representative images of *Emi1*-infected cells after EdU incorporation (photos after 96h post-infection) or immunostained for mitosis figures (with H3S10p) (photos after 24h post-infection). Centriole staining by FOP is used to identify progenitors (black arrows) and differentiating deuterosomal cells (white arrows). **(C)** Proportion of differentiating deuterosomal cells amongst all the cells marked by EdU after 96h post-infection by *GFP*, *Ccne2* or *Ccne2*+*Emi1* (left), of amongst the cells harboring mitosis figures 24h post-infection by *GFP*, *Ccna2* or *Ccna2*+*Emi1* (right). **(D)** Representative images of *Ccne2*+*Emi1* co-infected (top) or *Ccna2+Emi1* co-infected (bottom) cells, after EdU incorporation (photos after 96h post-infection) or immunostained for phosphorylated H3 serine 10 (H3S10p) (photos after 24h post-infection). Centriole staining by FOP is used to identify progenitors (black arrows) and differentiating deuterosomal cells (white arrows).

Cyclin E2 and A2 are two other cell cycle core regulators not re-activated during MCC differentiation. Cyclin E2 is required for the transition between G1 and S and the initiation of replication whereas Cyclin A2 is crucial for the correct progression through S and G2 phases up to mitosis entry (*27*, *28*). We infected cells with Cyclin E2 and Cyclin A2 and monitored DNA replication and mitosis entry events, respectively. We observed that Cyclin E2 infection alone was sufficient to partially induce DNA replication in differentiating cells – marked by foci of incorporated EdU 96hpi (**Fig. 5C**, left, **5D**) – whereas Cyclin A2 infection alone was sufficient to induce pseudo-mitosis – marked by nuclear envelope breakdown and chromosome condensation (**Fig. 5C**, right, **5D, S9**).

Importantly, during the cell cycle, EMI1 inhibits APC/C complex and stabilizes cyclins in S and G2 phases (*25*). When infecting cell cultures with a combination of Cyclin E2 or Cyclin A2 and *Emi1*, we observed a synergistic increase of the proportion of differentiating cells incorporating EdU or entering into pseudo-mitosis, respectively (**Fig. 5C**). Interestingly, full DNA replication and cytokinesis events were never observed, suggesting that even though replication and cytokinesis factors are widely co-opted in MCC (**Fig. S4**), some key factors are missing in addition to cyclins and APC/C regulators.

These results show that re-expressing *Emi1*, *Ccne2* or *Ccna2* in differentiating MCCs is sufficient to partially rescue DNA replication and mitosis entry. They further outline the synergy between EMI1 and these cyclins which can be recapitulated during this cell cycle variant. Altogether, these results confirm that MCC differentiation is a genuine cell cycle variant and show that it is tailored by variation in cyclin and APC/C inhibitor expression to allow centriole biogenesis without associated nuclear replication and mitosis. In a follow-up study, we investigate the role of Cyclin O, the first non-canonical cyclin expressed during the MCC cell cycle variant (**Khoury Damaa et al., 2024, ref. 44**).

## 4 Discussion

The cell cycle machinery represents an evolutionary conserved gearset crucial for the regulation and progression of cell division which involves DNA replication and mitosis. Some variants of the cell cycle have been shown to exist, e.g. cells can skip DNA replication to make haploid gametes in meiosis or to accelerate cell division during asynthetic fission (*29*). In other variants, cells skip mitotic cell division to increase genomic content of differentiating cells (endoreplication; (*30*)). Here, we show that differentiation of MCC is a final and customized iteration of the cell cycle skipping both DNA replication and mitotic division, to drive the amplification of cytoplasmic organelles, the centrioles (**Fig. 6**). We propose this process to be a genuine variation of the cell cycle: (1) cells within the MCC cell cycle variant are organized in a circular transcriptomic trajectory; (2) cells with S, G2 and M-like signatures are sequentially distributed along this circular trajectory; (3) they progressively increase their transcriptional output, followed by a sharp decline when cells are in G2/M-like phase; (4) waves of canonical and non-canonical cyclins demarcate consecutive functional phases along this circular trajectory; (5) these cyclins play a predominant role in regulating the progression of differentiation and (6) re-expression of missing core cell cycle partners can restore partial DNA replication and mitosis.

**Fig. 6:**
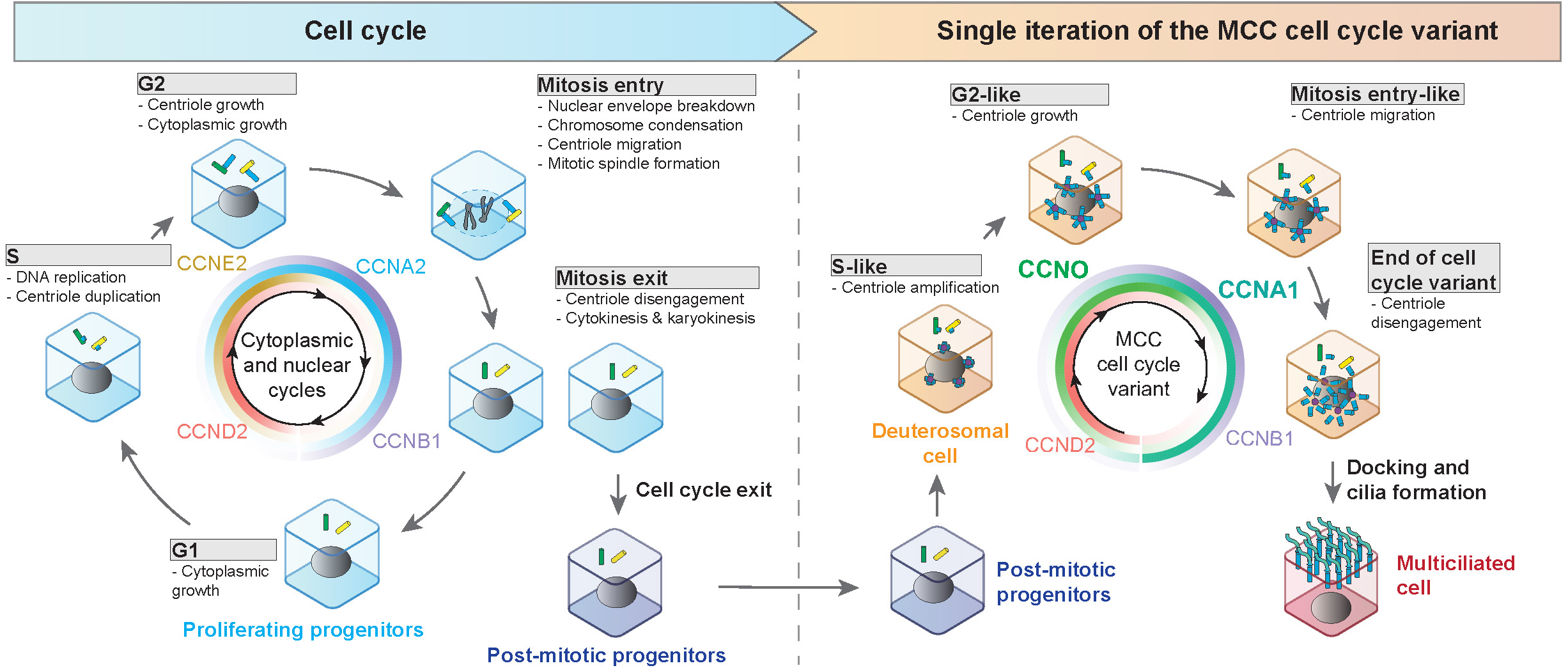
Model of deuterosomal cells undergoing a final iteration of a cell cycle variant during differentiation into MCCs. Canonical cyclins orchestrate proliferation of radial glial progenitor cells. Eventually, progenitors exit the cell cycle and commit to an ependymal fate. A final iteration of a cell cycle variant occurs to sustain centriole amplification, delineated by successive waves of canonical and non-canonical cyclins. The expression of each cyclin demarcates “pseudo” cell cycle phases, in which the transcriptional signature resembles that of cycling cells.

Cells undergoing endoreplication differ from dividing cells by the absence of their ability to segregate chromosomes or divide. This is partly driven by a tuning of CDK activity allowing the cell to progress through S but not through M-phase (*30*). Here we show that the core repressor of APC/C, EMI1, is missing in the MCC cell cycle variant, and that its re-expression is sufficient to trigger partial DNA replication and mitotic events. Correspondingly, a previous study focusing on differentiating skin MCCs in Xenopus, showed that the overexpression of *Emi2*, an *Emi1* homolog, also leads to DNA replication and mitotic events (*5*). We propose that, in the absence of strongly expressed APC/C inhibitors, an overactive APC/C dampens the MCC cell cycle variant even more than in endoreplication, effectively skipping DNA replication (**Fig. S10A**). This would eventually lead to an entire decoupling between the nuclear and cytoplasmic compartments, as seen in developing fly blastoderms (*31*). Since decoupling between centrosome and nuclear cycles can also be obtained by dampening CDK activity in cycling cells (*32*, *33*) and that DNA replication and mitotic events can be restored in cells undergoing this MCC cell cycle variant by disinhibiting CDK activity (*3*, *5*), we suggest the existence of a cytoplasmic -or centriolar-CDK threshold, lower than the S-phase threshold. A quantitative control of CDK activity by APC/C and the regulated expression of its inhibitors *Emi1/2* would thus allow the progression of a purely cytoplasmic MCC cell cycle variant.

We highlight that while the core cell cycle machinery is present, the E2 and A2 cyclins are respectively replaced by the non-canonical O and A1 cyclins. Rescuing the activity of the canonical cyclins E2 and A2 is sufficient to partially reroute differentiating cells towards DNA replication or chromosome condensation. This shows that a qualitative control, in addition to the APC/C-mediated quantitative regulation of CDK1 activity, orchestrates centriole amplification rather than DNA replication and mitosis in MCC (**Fig. S10B**). Of note, the expression of these cyclins in two distinct waves with Cyclin O ahead of Cyclin A1 contrasts with what has been suggested during MCC differentiation in respiratory cells (*4*). This could be explained by the great increase in the resolution of the progression of differentiation achieved by scRNAseq in our study. In conclusion, a nearly identical cell cycle machinery with identical transcriptional oscillations but different sets of cyclins orchestrates the MCC cell cycle variant, which is entirely different from cell division. In a follow-up study, we have further characterized the role of the non canonical Cyclin O in this MCC cell cycle variant (**Khoury Damaa et al., 2024, ref. 44**) and reveal that it is required to progress through the G1-to-S transition of this cell cycle variant, thereby reinforcing the similarities with regulatory mechanisms used in the canonical cell cycle.

Importantly, the successive expression of non-canonical cyclins O and A1, as well as the tuned expression of the *Emi2*, replacing *Emi1* to regulate APC/C activity, are also observed in male meiosis, an important variant of the canonical cell cycle where centrioles are produced, uncoupled from DNA replication (*34–36*). The wave-like expression of *Ccno* is also associated with the transcriptional silencing of *Ccne2* during male meiosis, further reinforcing the relevance of the MCC differentiation program as a genuine cell cycle variant, regulated both qualitatively and quantitatively.

While such proximity between cell division and centriole amplification may appear risky (*1*), this cell cycle variant – marked by the association of centriole amplification with low CDK activity – could be an efficient barrier to avoid DNA replication and cell division in cells with amplified centrioles. Supporting this “fail-safe” hypothesis, several medical reports reveal the existence of MCC in cysts that have been found in a wide variety of normally non multiciliated-organs (e.g. in (*37–40*)). Such potential for multiciliation in a progenitor cell could also underpin the observed formation of MCC along the lumen of non-MCC fluid producing tissues (kidney, urethra) in pathological situations where they could contribute to restore fluid flow (*41–43*). In addition to providing a detailed prism for the study of multiciliation mechanisms, the close similarity of multiciliated differentiation mechanisms to those of the cell cycle revealed in this study expands the putative functions of multiciliated cells, sheds new light on the link between centriole number and cell division, and reveals that cell cycle variants can also control cytoplasmic and not only nuclear processes during cell differentiation.

## Supporting information

Supplemental informations

## Acknowledgments

We thank all members of the Spassky and the Koszul labs for fruitful discussions and comments on the manuscript, Andrew Holland and Damien Coudreuse for reading and discussing the manuscript. We are grateful to Kévin Lebrigand and Virginie Magnone for fruitful discussions on single cell RNA sequencing, A.-K. Konate for administrative support, and the IBENS Animal Facility for animal care.

## Author contributions

Conceptualization: J.S., R.K., A.M.; Methodology: J.S., A.-R.B., M.F., N.D.; Software: J.S.;

Formal analysis: J.S.; Investigation: J.S., A.-R.B., M.K.D., R.B., L.-E.Z., N.S., A.M.; Resources:

A.-R.B., M.K.D., M.F., N.D., N.S., A.M.; Visualization: J.S.; Funding acquisition: P.B., N.S., R.K., A.M.; Project administration: R.K., A.M.; Supervision: R.K., A.M.; Writing – original draft: J.S., R.K., A.M.

## Data and material availability

All raw and processed sequencing data generated in this study have been submitted to the NCBI Gene Expression Omnibus (GEO; https://www.ncbi.nlm.nih.gov/geo/) and will be publicly released upon publication.

