## Supplemental informations for "Identification of a new cell cycle variant during multiciliated cell differentiation"

**This file contains :**
**Figure S1 to S10 + legends**
**Legend Table S1**
**Material and Methods**

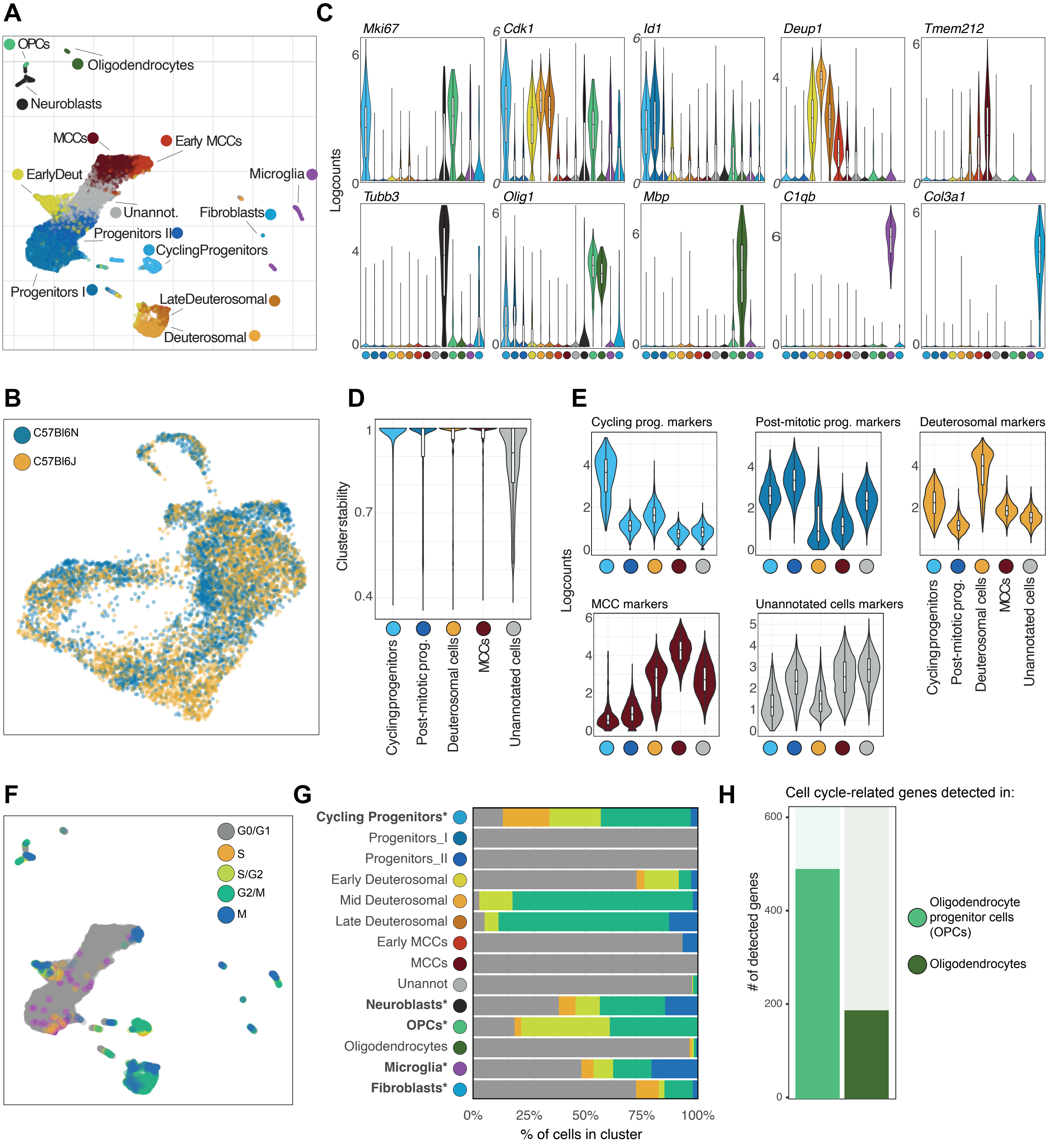


Fig S1 - Serizay et al.

**Fig. S1: Single-cell profiling of in vitro differentiating mouse radial glial cells into multiciliated cells**

**(A)** Single-cell RNA-sequencing profiling of in vitro differentiating mouse radial glial cells into multiciliated cells. All 16,401 cells are shown in this UMAP projection.

**(B)** UMAP projection of cells undergoing MCC differentiation. 4,000 randomly sampled cells from each replicate are shown. Colors represent the replicates.

**(C)** Expression of known markers of progenitors, differentiating or terminally differentiated MCCs, neuroblasts, OPCs and oligodendrocytes, microglia or stromal cells.

**(D)** Stability of cell population clusters. Note that the Unannotated cells have a low cluster stability, indicating that these cells do not reliably form a specific cluster.

**(E)** Expression of the top 5 markers of each group of cells, across the five main groups of cells.

**(F)** Putative cell cycle phase annotations inferred using SingleR using neural stem cell reference (*7*). All 16,401 cells are shown in this projection.

**(G)** Distribution of putative cell cycle phase annotations within each cell population. Cell populations known to proliferate are indicated with an asterisk.

**(H)** Number of cell cycle related factors detected in oligodendrocyte progenitor cells (OPCs) and post-mitotic oligodendrocytes (similar to Fig. 1D). A gene is considered expressed in an individual cell population if its expression is greater than 0.5 (logcounts) in at least 20% of the cells within the cell population.

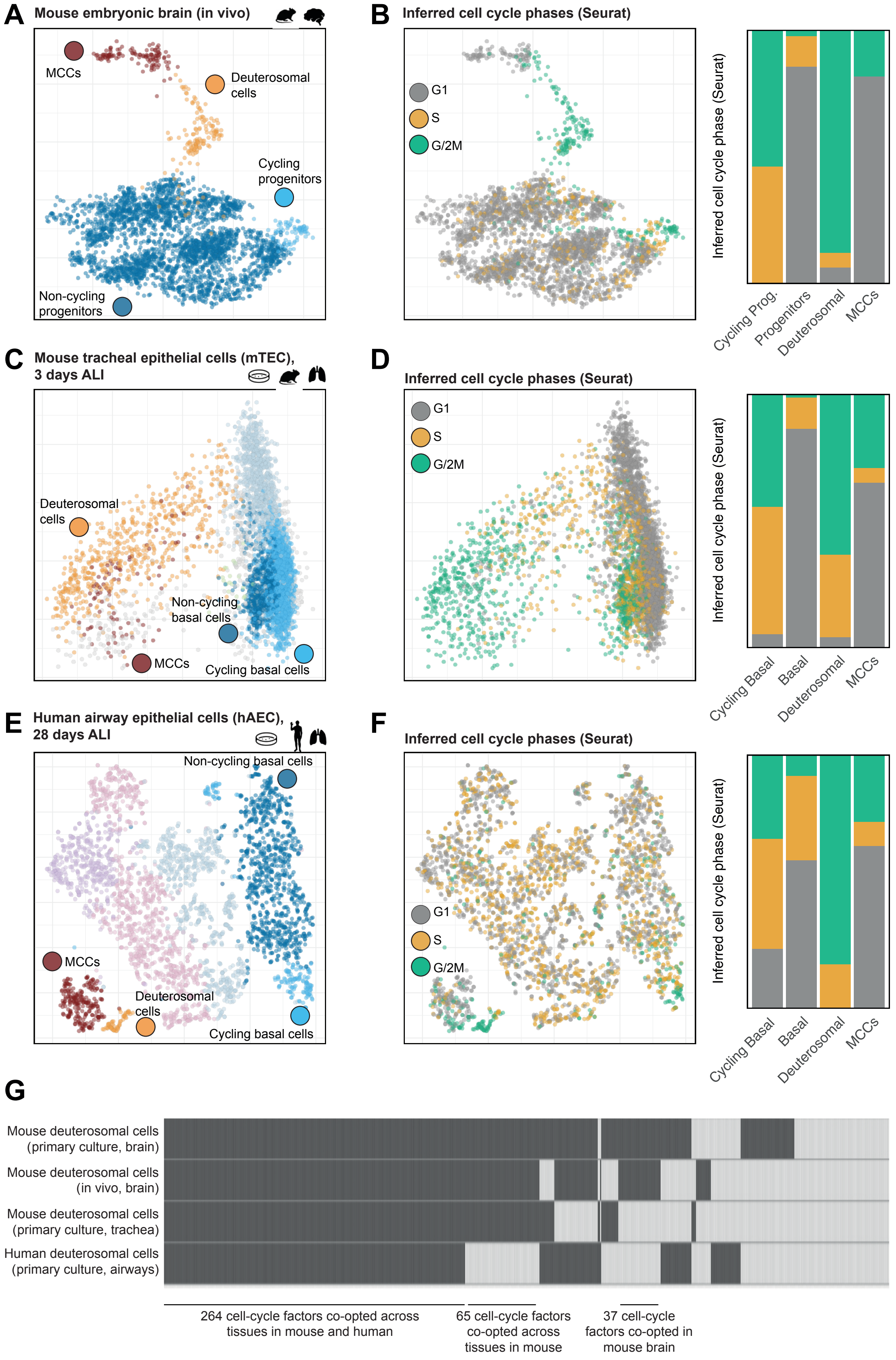


### Fig S2 - Serizay et al.

**Fig. S2: Cell cycle factors are reused for multiciliation across mammals**

- **(A)** UMAP embedding of individual transcriptomes from an in vivo scRNAseq experiment performed in brains from mouse embryos (La Manno et al., 2021). Color code indicates the different cell groups.
- **(B)** UMAP embedding of the in vivo brain scRNAseq dataset, with color code indicating the putative cell cycle phase annotations inferred using Seurat (left). Distribution of the putative cell cycle phase annotations in each cell group (right).
- **(C)** UMAP embedding of individual transcriptomes from an in vitro scRNAseq experiment performed in an air-liquid interface culture of mouse tracheal epithelial cells (mTEC) (Ruiz Garcia et al., 2019). Color code indicates the different cell groups.
- **(D)** UMAP embedding of the in vitro mTEC scRNAseq dataset, with color code indicating the putative cell cycle phase annotations inferred using Seurat (left). Distribution of the putative cell cycle phase annotations in each cell group (right).
- **(E)** UMAP embedding of individual transcriptomes from an in vitro scRNAseq experiment performed in an air-liquid interface culture of human airway epithelial cells (hAEC) (Ruiz Garcia et al., 2019). Color code indicates the different cell groups.
- **(F)** UMAP embedding of the in vitro hAEC scRNAseq dataset, with color code indicating the putative cell cycle phase annotations inferred using Seurat (left). Distribution of the putative cell cycle phase annotations in each cell group (right).
- **(G)** Summary of cell cycle related factors co-opted in in vitro mouse brain scRNAseq, in vivo mouse brain scRNAseq, in vitro mouse tracheal scRNAseq and in vitro human airway scRNAseq.

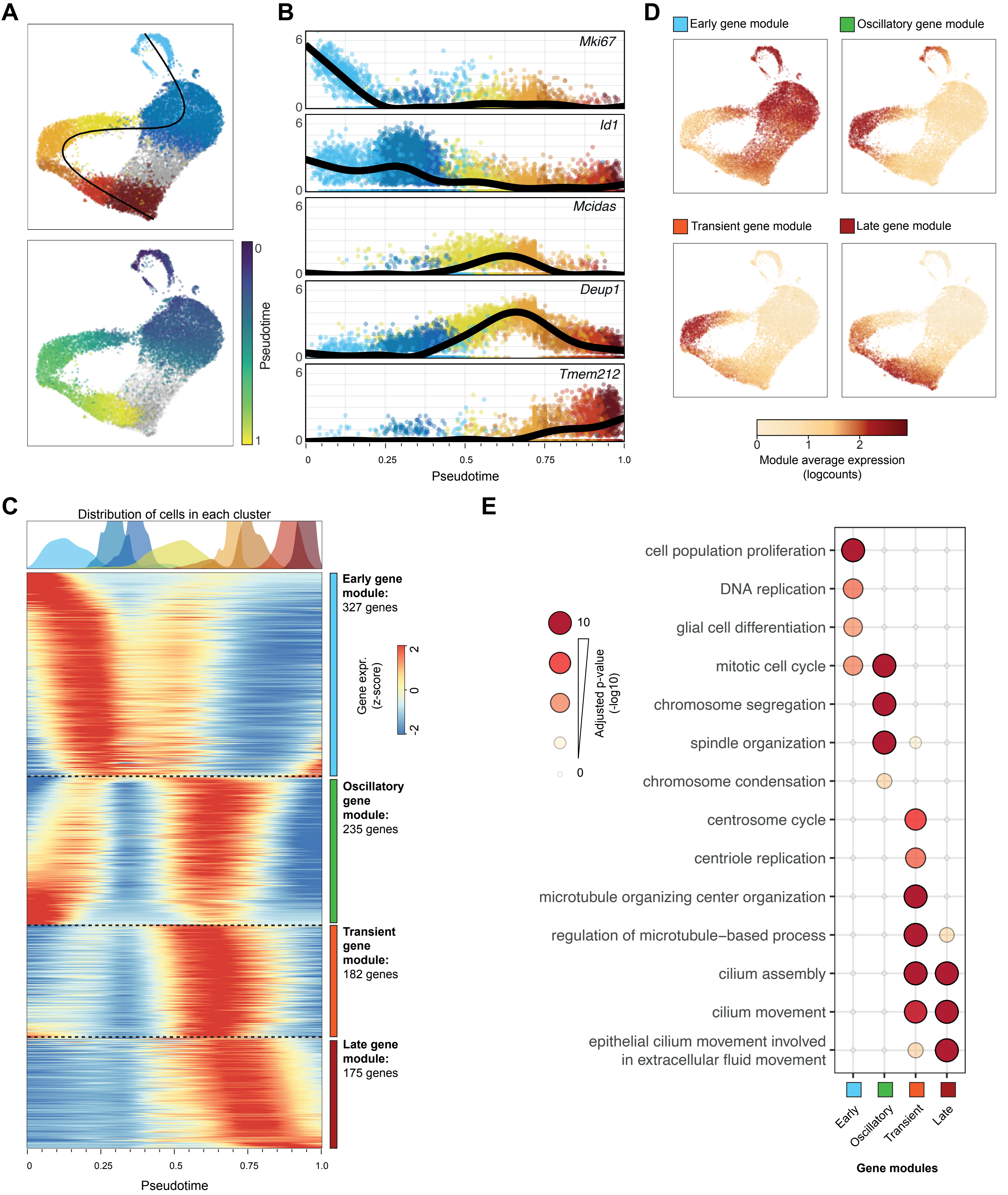


**Fig S3 - Serizay et al.**

**Fig. S3: Lineage of in vitro differentiating mouse radial glial cells**

- **(A)** Trajectory analysis of in vitro differentiating mouse radial glial cells into multiciliated cells. The cell lineage starts with cycling progenitors, passes through post-mitotic progenitors, deuterosomal cells and finally reaches MCCs. Top: cell lineage embedded in UMAP cell projection. Bottom: UMAP cell projection with cells colored by their inferred pseudotime (rescaled between 0 and 1).
- **(B)** Temporal expression of Mki67, Id1, Mcidas, Deup1 and Tmem212 along the differentiation trajectory shown in A.
- **(C)** Aggregated temporal expression of the 919 genes differentially expressed in the scRNAseq dataset. Color scale represents the gene expression (z-scored for each gene independently). On top of the heatmap, the distribution of cells in each cluster is shown. We distinguished four broad patterns of variation of gene expression amongst the 919 genes differentially expressed between cell clusters: a module of "early genes" expressed in progenitors and early deuterosomal cells, a module of "oscillatory genes" (highly expressed in cycling progenitors then silenced in non-cycling progenitors and reactivated in differentiating progenitors), a module of "transient genes" expressed only in deuterosomal cells and a module of "terminal genes" expressed in differentiated MCCs.
- **(D)** Average expression of each gene module in the UMAP cell projection.
- **(E)** Gene ontology over-representation analysis for each module depicted in C. This confirmed that the "oscillatory genes" module was enriched for cell cycle-related processes.

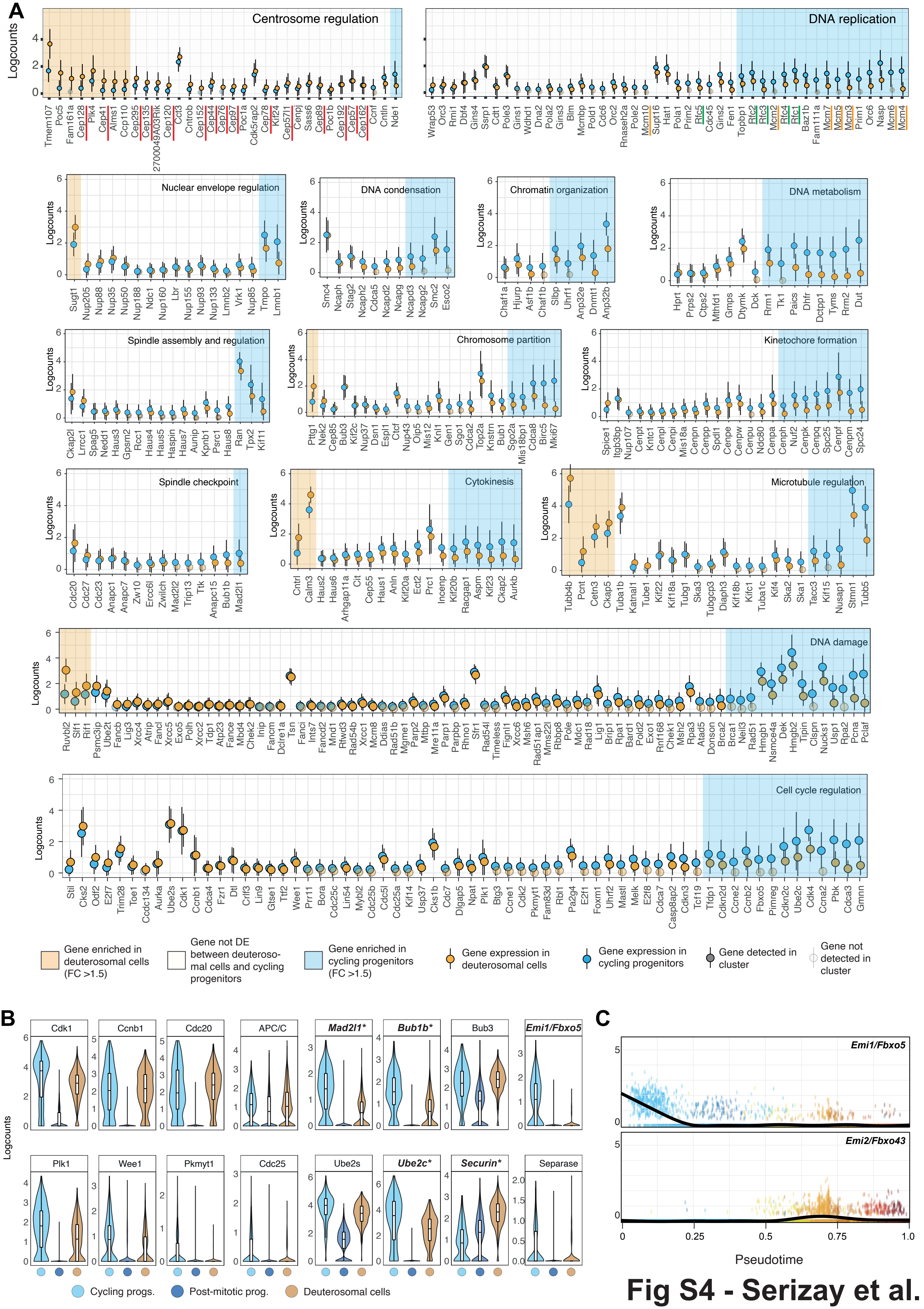


Fig S4 - Serizay et al.

**Fig. S4: Factors from all cell cycle subprocesses are re-expressed in deuterosomal cells**

**(A)** Expression of cell cycle factors involved in different cell cycle subprocesses, in deuterosomal cells (orange point+ranges) and in cycling progenitors (blue point+ranges). Transparent points denote genes not detected in one or the other cell population. Genes which are not detected in any of the two cell populations are not shown.

**(B)** Violin plot of the expression of the main components of the mitotic oscillator in different cell clusters. Labels in bold indicate factors differentially expressed between deuterosomal cells and G2/M cycling progenitors.

**(C)** Temporal expression of Emi1 and Emi2 along the differentiation trajectory shown in Fig. S3A.

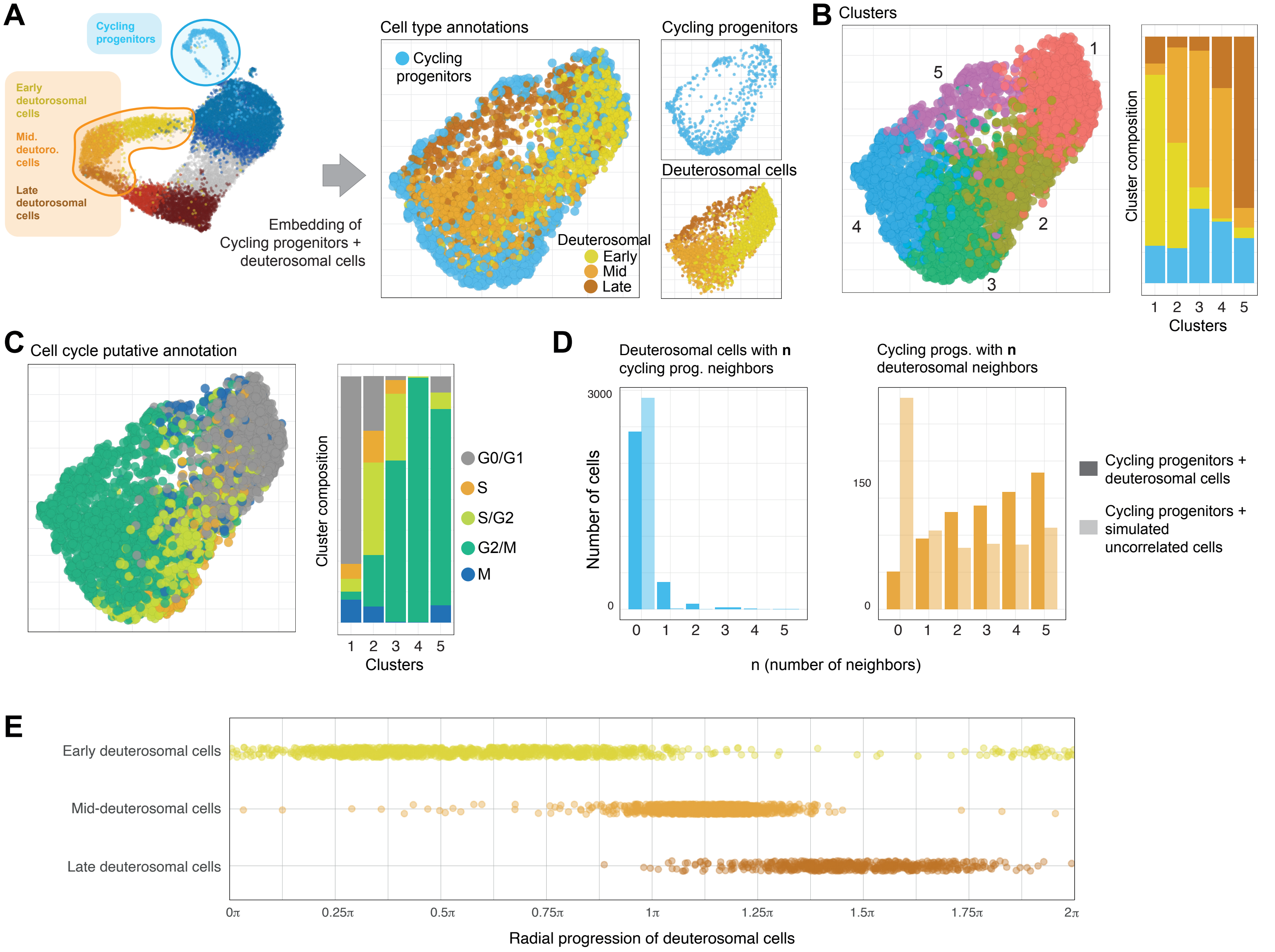


### Fig S5 - Serizay et al.

**Fig. S5: Sub-processing of cycling progenitors and deuterosomal cells**

**(A)** Shared UMAP embedding of all deuterosomal cells and all cycling progenitor cells, showing that cycling progenitors and deuterosomal cells aggregate in close proximity to each other in lower dimensional space.

**(B)** Shared cell clustering of all deuterosomal cells and all cycling progenitor cells. Right: proportion of each cell type per cluster. Each of the newly inferred clusters contain a mixture of roughly 15-20% of cycling progenitors and 80-85% of deuterosomal cells.

**(C)** Putative cell cycle phase annotations inferred using SingleR using neural stem cell reference (*7*). Deuterosomal cells and cycling progenitor cells are plotted in their shared UMAP embedding. Cycling progenitors and deuterosomal cells in each cluster are enriched for specific cell cycle phases.

**(D)** Number of cycling progenitor cells amongst the 5 nearest neighbors for each deuterosomal cell (left) and number of deuterosomal cells amongst the 5 nearest neighbors for each cycling progenitor cell (right). The observed distribution of cycling progenitors / deuterosomal cells amongst the 5 nearest neighbors is shown in solid colors, while the same distribution measured between real cycling progenitors and simulated, uncorrelated cells is shown in transparent.

**(E)** Distribution of early, mid- and late deuterosomal cells along their radial progression.

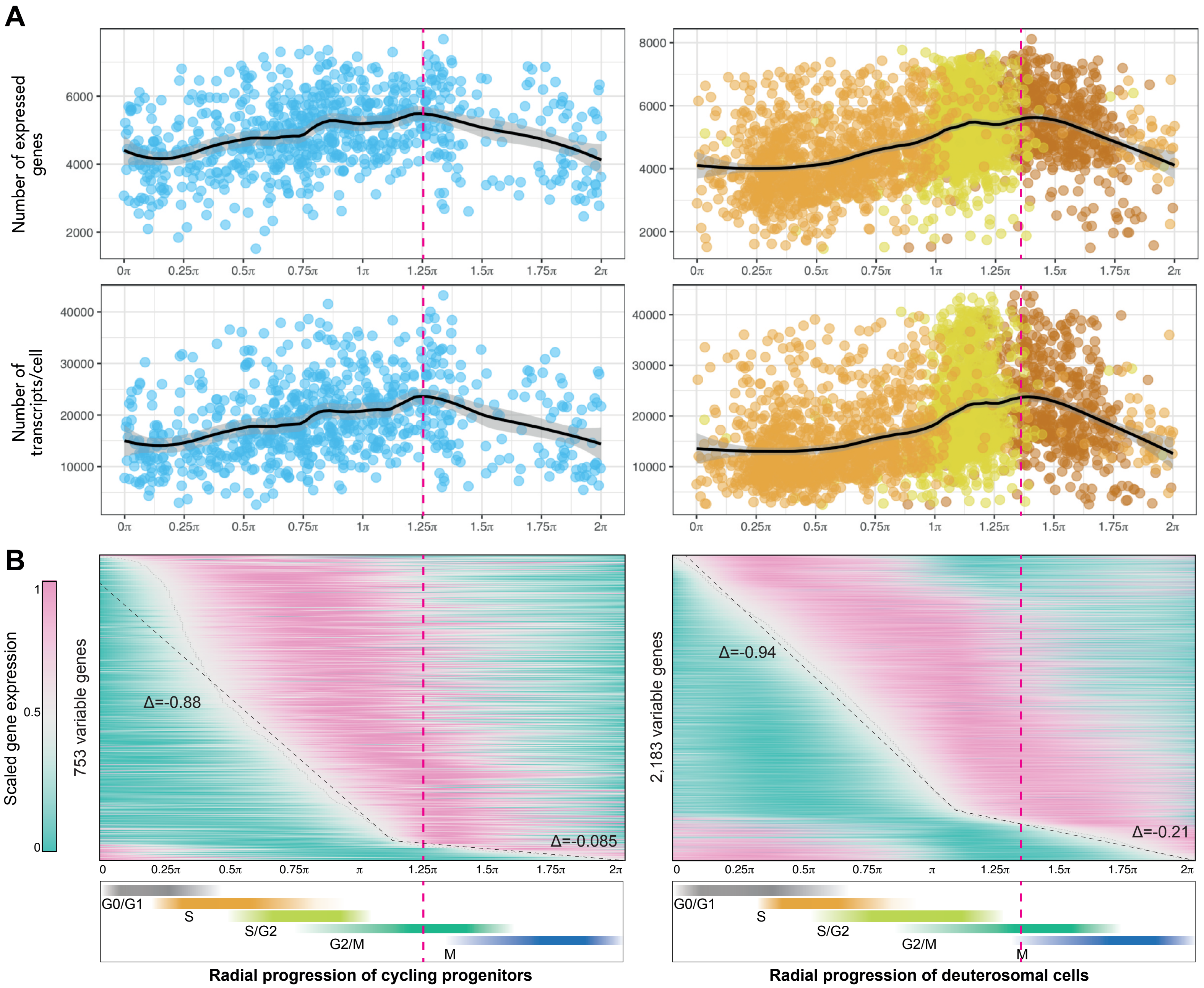


Fig S6 - Serizay et al.

**Fig. S6: Deuterosomal cells share features of transcriptional dynamics from cycling cells**

**(A)** Number of expressed genes (top) and of transcripts per cells (bottom), in cycling progenitor cells (left) or in deuterosomal cells (right) distributed according to their radial progression.

**(B)** Temporal expression of genes which expression vary in cycling progenitor cells (left) or in deuterosomal cells (right). Expression is normalized to the interval [0;1] for each gene, and genes are ordered by the time when 0.5 is crossed from below (seen as a white line). The slope of the white line reports the rate of transcription onsets per unit time. The steeper the slope, the higher the rate is. The rate of transcription onsets is generally constant in cycling progenitor cells and in deuterosomal cells until ~ 1.20π. At this time point, the rate of transcription onsets dramatically decreases (10-fold in cycling progenitors and 4.5-fold in deuterosomal cells).

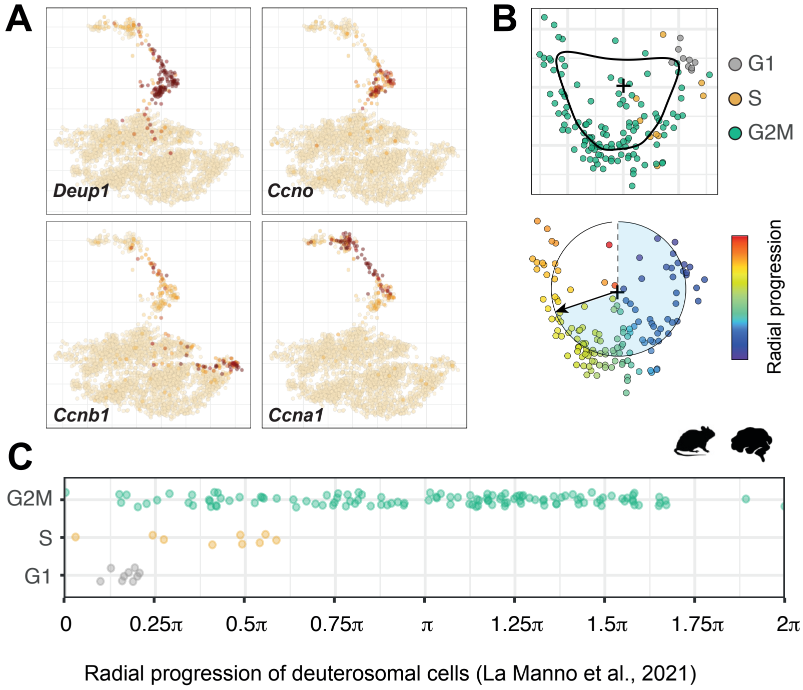


**Fig S7 - Serizay et al.**

**Fig. S7: Cell cycle factors are reused for multiciliation across mammals**

**(A)** Expression of Deup1 and three cyclins in the in vivo brain scRNAseq dataset (15).

**(B)** Top: PCA embedding (PC1 and PC2) of deuterosomal cells only. The black curve denotes the average position of cells around the center point of the PCA space. Bottom: representation of the deuterosomal cells with a color range indicating the radial progression of each cell.

**(C)** Radial distribution of deuterosomal cells with a color scale indicating the putative cell cycle phase annotated using Seurat.

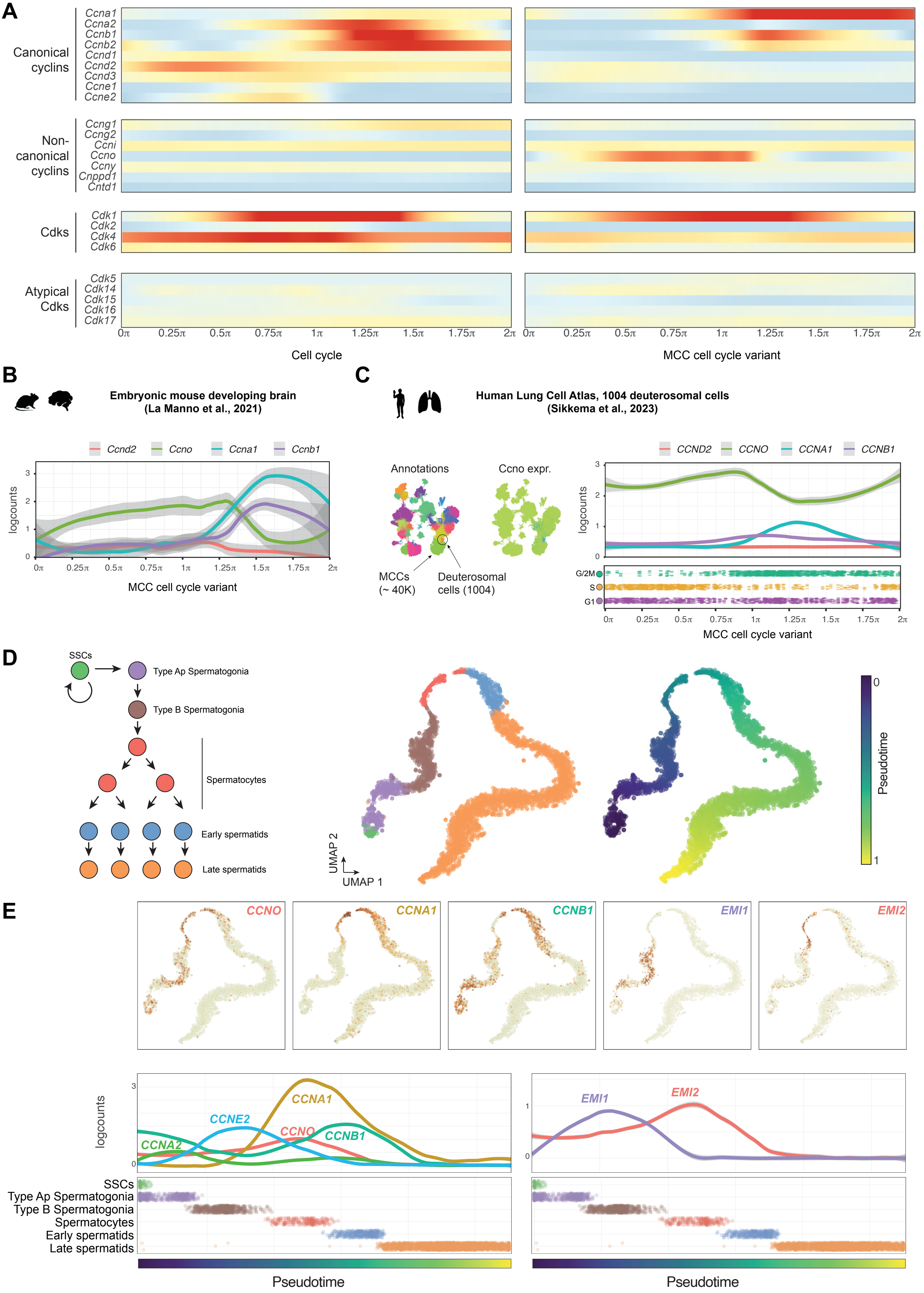
A
Canonical cyclins
Ccna1
Ccna2
Ccnb1
Ccnb2
Ccnd1
Ccnd2
Ccnd3
Ccne1
Ccne2
Non-canonical cyclins
Ccng1
Ccng2
Ccni
Ccno
Ccny
Cnppd1
Cntd1
Cdks
Cdk1
Cdk2
Cdk4
Cdk6
Atypical Cdks
Cdk5
Cdk14
Cdk15
Cdk16
Cdk17
Cell cycle
MCC cell cycle variant
B
Embryonic mouse developing brain (La Manno et al., 2021)
Ccnd2
Ccno
Ccna1
Ccnb1
logcounts
MCC cell cycle variant
C
Human Lung Cell Atlas, 1004 deuterosomal cells (Sikkema et al., 2023)
Annotations
Ccno expr.
MCCs (~ 40K)
Deuterosomal cells (1004)
CCND2
CCNO
CCNA1
CCNB1
logcounts
G/2M
S
G1
MCC cell cycle variant
D
SSCs
Type Ap Spermatogonia
Type B Spermatogonia
Spermatocytes
Early spermatids
Late spermatids
UMAP 2
UMAP 1
Pseudotime
E
CCNO
CCNA1
CCNB1
EMI1
EMI2
logcounts
CCNA1
CCNE2
CCNO
CCNB1
CCNA2
EMI1
EMI2
SSCs
Type Ap Spermatogonia
Type B Spermatogonia
Spermatocytes
Early spermatids
Late spermatids
Pseudotime
Pseudotime

**Fig. S8: Expression of cyclins during ciliated cell differentiation**

- **(A)** Temporal expression of canonical and atypical cyclins along the cell cycle (left) or along the MCC cell cycle variant (right).
- **(B)** Expression of Cyclin D2 (Ccnd2), Cyclin O (Ccno), Cyclin A1 (Ccna1) and Cyclin B1 (Ccnb1) cyclins in deuterosomal cells distributed according to their radial progression, in in deuterosomal cells from the embryonic mouse developing brain (La Manno et al., 2021).
- **(C)** Expression of Cyclin D2 (CCND2), Cyclin O (CCNO), Cyclin A1 (CCNA1) and Cyclin B1 (CCNB1) cyclins in deuterosomal cells distributed according to their radial progression, in deuterosomal cells from the adult human airway epithelium (Sikkema et al., 2023).
- **(D)** Embedding of a human spermatogenesis scRNAseq dataset (Guo et al., 2018) in UMAP 2D dimensional space. Cells are colored by their cell type (middle) or by the pseudotime inferred with slingshot (right). SCCs: Spermatogonial stem cells.
- **(E)** Expression of Cyclin O (CCNO), Cyclin A1 (CCNA1) and Cyclin B1 (CCNB1) cyclins during spermatogenesis, in UMAP projection (**A**) or along the inferred pseudotime (**B**).

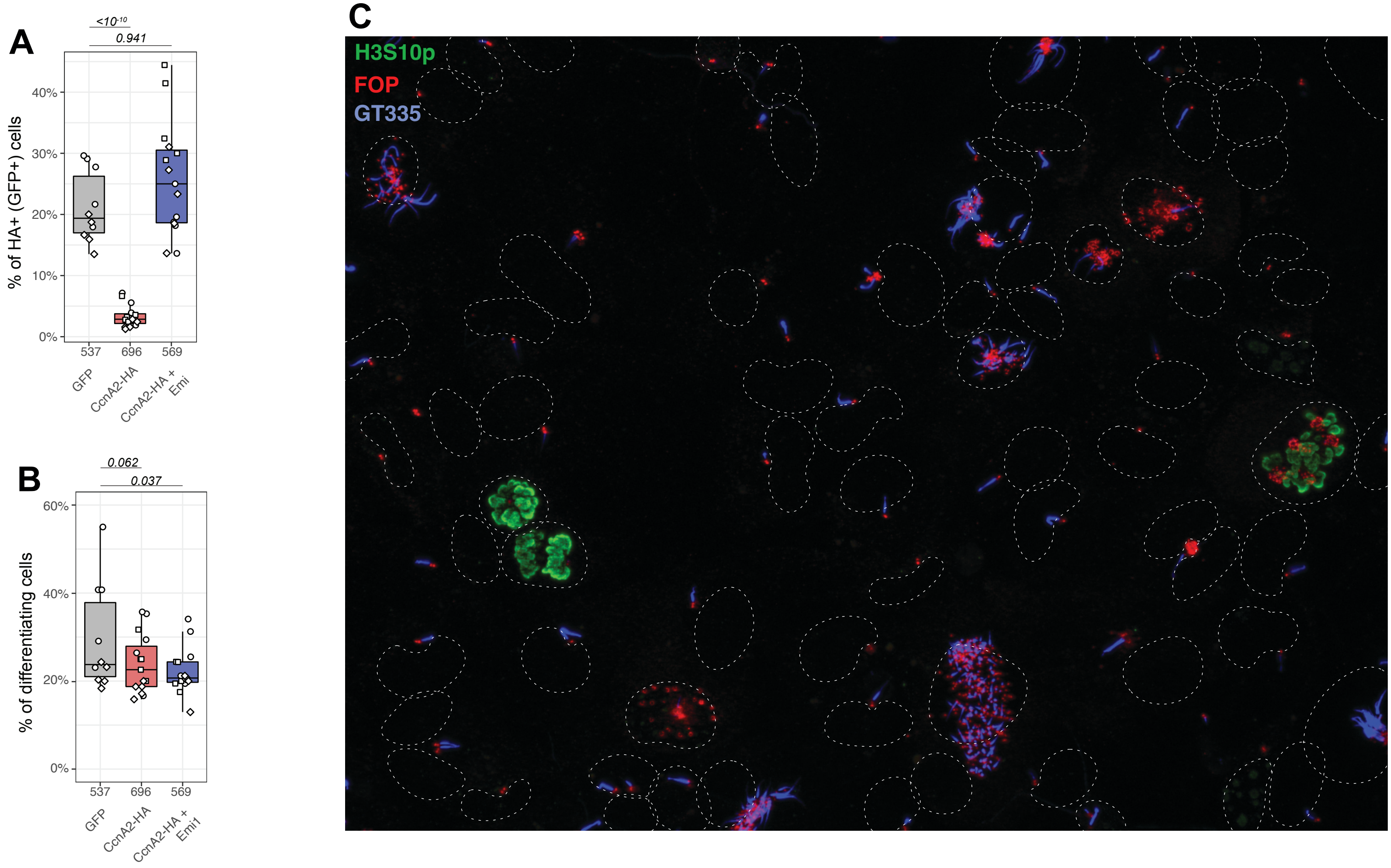


Fig S9 - Serizay et al.

**Fig. S9: Canonical Cyclin A2 expression in differentiating MCCs leads to abnormal mitosis figures**

**(A)** Proportion of HA+ (or GFP) cells in WT radial glial cell cultures infected by GFP, CcnA2-HA or CcnA2-HA+Emi1-Myc. The total number of cells counted across three biological replicates is indicated for each experiment. P-values from batch-corrected t-tests are displayed on top of the graph.

**(B)** Proportion of differentiating cells in WT radial glial cell cultures infected by GFP, CcnA2-HA or CcnA2-HA+Emi1-Myc. The total number of cells counted across three biological replicates is indicated for each experiment. P-values from batch-corrected t-tests are displayed on top of the graph.

**(C)** Representative immunofluorescence image of an in vitro cell differentiation culture infected with CcnA2-HA+Emi1-Myc and labeled with pH3 to mark mitosis figures and FOP to mark centrioles. Two different types of mitosis figures are observed in the same field (40X): left: two normal cycling progenitors (with only 2 pairs of duplicated centrioles each) are dividing, as marked by H3S10p; right: an abnormal differentiating MCC (with several tens of amplified centrioles) exhibits condensed chromosomes marked by H3S10p.

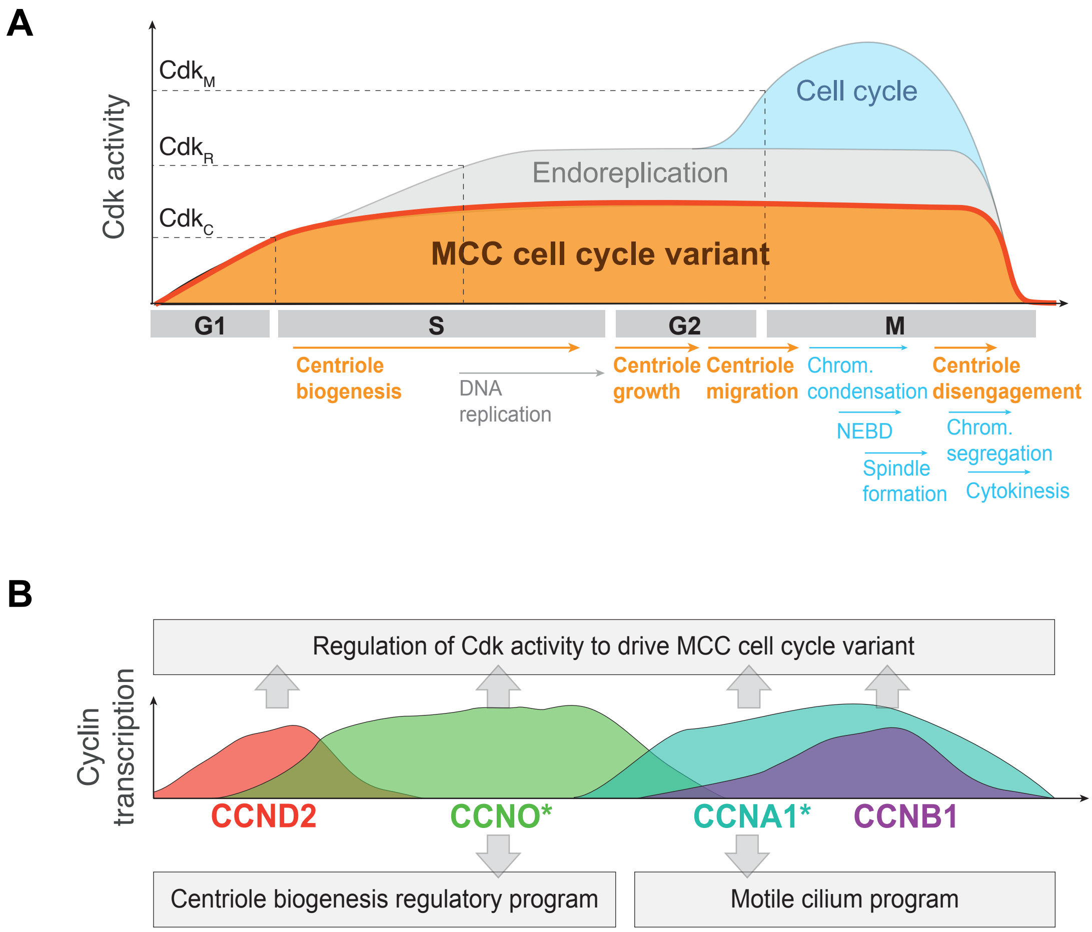


**Fig S10 - Serizay et al.**

**Fig. S10: Cdk activity and cyclin roles during the MCC cell cycle variants**

**(A)** Model of regulated Cdk activity during the MCC cell cycle variant. Three different thresholds of Cdk activity are represented. We propose that in differentiating MCCs, the overall Cdk activity reaches the lowest threshold ($Cdk_C$), where centriole biogenesis and potentially other cell cycle-like cytoplasmic reorganization processes are regulated. The overall Cdk activity however never reaches the replication threshold ($Cdk_R$) described in endoreplicating cells, and thus never triggers DNA replication. The $Cdk_M$ threshold, described to be above the $Cdk_R$ threshold, is therefore never attained. $Cdk_R$ and $Cdk_M$ thresholds can however be attained in MCC under Cdk disinhibition.

**(B)** Sequential waves of cyclin expression during MCC cell cycle variant are represented, along with the transcriptional programs synchronized with the expression of Ccno and Ccna1.

**Table S1: Average and median gene expression in single-cell RNA-seq of in vitro differentiating multiciliated cells**

Cells are aggregated by their cluster annotation and the average expression across each cluster is calculated for each gene. Within each cluster, the proportion of cells expressing each gene is indicated (using a minimum threshold of 2 counts).

#### Materials and Methods

##### Single-cell RNA-seq of in vitro differentiating multiciliated cells

Newborn mice (P0-P2) were sacrificed by decapitation. The brains were then dissected in Hank's solution (10% HBSS, 5% HEPES, 5% sodium bicarbonate, 1% penicillin/streptomycin (P/S) in pure water) and the extracted ventricular walls were cut manually into pieces, followed by enzymatic digestion (DMEM glutamax, 33% papain (Worthington 3126), 17% DNase at 10 mg/ml, 42% cysteine at 12 mg/ml) for 45 min at 37 °C in a humidified 5% CO2 incubator. Digestion was stopped by addition of a solution of trypsin inhibitors (Leibovitz Medium L15, 10% ovomucoid at 1 mg/ml, 2% DNase at 10 mg/ml). The cells were then washed in L15 and resuspended in DMEM glutamax supplemented with 10% fetal bovine serum (FBS) and 1% P/S in a Poly-L-lysine (PLL)-coated flask (equivalent of 2 ventricular walls per 25cm2 flask). Ependymal progenitors proliferated for 4-5 days until confluence, followed by shaking (250rpm) overnight to remove contaminant cells (neurons, oligodendrocytes, etc.). The day after shaking, the flasks' medium was changed to warm serum-free DMEM glutamax 1% P/S to trigger ependymal cell differentiation for 2 days. At this time point, cells at all stages of cell cycle or ependymal differentiation are present in the flask. After 2 days of in vitro differentiation, flasks were rinsed with PBS 1X twice and treated by enzymatic cell dissociation (Trypsin, 1mL) for 10 min and trituration to obtain a single cell suspension. Digestion was stopped by addition of 1 ml of fetal bovine serum (FBS). The cells were then washed in serum-free DMEM glutamax 1% P/S and resuspended in HBSS-0.1% BSA for single-cell RNA-seq library preparation. Cell cultures from three different animals with the same genetic background and treatment were pooled together; to multiplex them, samples were labelled using a cell surface protein labelling strategy following manufacturer's instructions (https://assets.ctfassets.net/an68im79xiti/5KA1NbZdTOam8A0yq6KyC2/f3e0479ff7b1c1633e6ddf8959a88a3b/CG000149_DemonstratedProtocol_CellSurfaceProteinLabeling_Rev_A.pdf), using TotalSeq® hashtag antibodies (BioLegend). However, the efficiency of the cell hashing did not allow to unambiguously separate cells coming from different animals and was therefore not used in downstream analysis. Cell suspensions were passed through a 40 µm Flowmi cell strainer (BelArt) and cell concentrations were carefully evaluated with a Countess FL automated cell counter (Thermofisher). For single cell RNA-seq, cells were partitioned with a Chromium equipment (10X Genomics) and libraries were prepared using the standard Single Cell 3' v3.1 protocol (10X Genomics). Sequencing was performed on an Illumina NextSeq 500 sequencing machine following manufacturer's instructions, on a high Flowcell, using the following sequencing cycles: 28b for read 1, 55b for read 2, 8b for index.

**Adenovirus infections**

The following adenovirus gene expression systems were ordered and packaged by VectorBuilder: pAV[Exp]-CMV>EGFP (control eGFP), pAV[Exp]-CMV>Myc/mFbxo5 (EMI1, tagged with myc in N-terminal, under the control of CMV), pAV[Exp]-mCherry-CMV>HA/mCcne2 and pAV[Exp]-mCherry-CMV>HA/mCcno (CCNE2 or CCNO, tagged with HA in N-terminal, under the control of CMV in a vector also expressing mCherry). Mouse Fbox5 (Emi1) mRNA (NM_025995.2), Ccne2 mRNA (NM_001037134.2) and Ccno mRNA (NM_001081062.2) references were used.

For adenovirus infections, ependymal cell cultures were performed as described above, although cells were plated on Poly-L-lysine (PLL)-coated 12mm coverslips in 24-well plates after tissue dissociation. After 2 days of in vitro differentiation (DIV2), adenovirus infections were performed by adding the adenovirus crude lysates to differentiating multiciliated progenitors (500 MOI for single infections, 500MOI each for co-infection with 2 viruses) for 4h, shaking every hour for optimal infection. After 4h of infection, cells were washed with pre-warmed 0.5% FBS-DMEM, and infected cells were kept in culture for 20h or 92h under serum-starved condition with 0% FBS-DMEM. The cells were then fixed and subjected to analysis according to days post infection (DPI).

##### Immunostainings and EdU incorporation stainings

EdU was added after 4h of viral infection, immediately after virus wash-out. Cell cultures were fixed 24h or 96h post-infection (20h or 90h EdU incorporation) in paraformaldehyde (PFA 4%) at 4°C for 10 minutes, then pre-blocked in saturation buffer (PBS 1X with 0.2% Triton X-100 and 10% FBS). Primary antibody immunostaining was performed for 1h at room temperature in the saturation buffer. When needed, EdU incorporation was labeled using the ThermoFisher Click-iT® EdU Imaging Kit (catalog # C10337) following manufacturer's recommendations. Cells were then washed in PBS and secondary antibody staining performed for 1h at room temperature in the saturation buffer. Cells were counterstained with DAPI (10 µg/ml, Sigma) for nucleus/DNA observation and mounted in DAPI Fluoromount (Southern Biotech). The following antibodies were used: rabbit anti-Histone 3 (pSer10; 1:100; 9701, Cell Signaling), mouse IgG2b anti-FOP (1:700), and species-specific Alexa Fluor® secondary antibodies (1:400; Invitrogen).

##### Computational analyses

Pre-processing of scRNAseq of in vitro differentiating multiciliated cells

Fastq file demultiplexing, barcode processing, gene counting, and aggregation were made using the Cell Ranger software following 10X Genomics guidelines. 10X Genomics mm10 genome reference and gene annotations compiled in 2020 were used (https://support.10xgenomics.com/single-cell-gene-expression/software/release-notes/build#mm10_2020A). Empty cells (detected by emtpyDrops from DropletUtils) were filtered out and only protein-coding genes were retained.

Analysis of scRNAseq of in vitro differentiating multiciliated cells

**Batch correction, dimensionality reduction, clustering:** Batch correction and replicate merging was performed using fast MNN correction from batchelor package, using only marker genes identified in either one of the replicates using scran package. For visualization, UMAP embedding was performed using 50 dimensions from the corrected PCA; the embedding was performed several times with a changing seed and a representative 2D projection of the dataset was used. Cell clustering was performed on the dataset embedded in a SNN graph (k=5 neighbors), using the Louvain algorithm from igraph.

**Cell annotation:** Clusters were manually annotated using cell markers identified in previous studies, and annotations were validated using automated annotation transfer (relying on SingleR package) from a reference collection of 358 bulk RNA-seq profiles of sorted cell populations.

**Gene annotation:** Genes in any given cell cluster were annotated as expressed as long as they had a log-normalized expression greater than 0.5 in more than 20% of the cells within the cell cluster. Genes differentially expressed between cell populations were identified using the "findMarkers" function from the scran package. Any gene with an expression fold-change greater than 1.5 between two cell clusters was considered enriched in the cluster with the greatest expression.

**Cluster stability:** Cluster stability was estimated by clustering 80% randomly sampled cells from the original dataset 30 times independently and comparing the clustering of each cell to the original clustering.

**Cell cycle phase annotation:** Putative cell cycle phases were transferred from a recent scRNAseq dataset of proliferating neural stem cells with annotated cell cycle phases (O'Connor et al., 2021) using SingleR. G0, G1 and Late G1 labels were collapsed to a single "G1" label.

**Trajectory analysis:** Trajectory and pseudotime inferences were performed using PCA embedded data with slingshot, specifying start (proliferating progenitors) and end clusters (terminally differentiated MCCs) to orientate the trajectory. Computed pseudotime was scaled between 0 and 1 for clarity. Continuous gene expression along the slingshot-inferred trajectory was modeled using a generalized additive model. For visualization of the continuous expression of many variable genes along the slingshot-inferred trajectory (e.g. in Fig. S2C), genes were first clustered using k-medoids then seriated using the spectral method from the seriation package.

**GO enrichment analysis:** GO over-representation analysis was conducted for individual gene sets using the gprofiler2 package.

**Combined analysis of cycling progenitors and deuterosomal cells:** Combined sub-processing of both cycling progenitor cells and all the deuterosomal cells was done by performing batch correction and replicate merging on the aggregated subsets of cells, using a large set of cell cycle-related genes obtained from GO mouse annotation database. For visualization, UMAP embedding was performed using 50 dimensions from the corrected PCA; the embedding was performed several times with a changing seed and a representative 2D projection of the dataset was used. Cell sub-clustering was performed using the same strategy as that used in the complete dataset (see hereinabove). The number of nearest cycling progenitor (deuterosomal) neighbors for deuterosomal cells (progenitors) (from zero to five) was calculated within the five nearest neighbors identified using Euclidean distance between pairs of cells embedded in corrected PCA space. This was performed either on the real cycling progenitors and deuterosomal merged dataset, or on a dataset containing real cycling progenitor cells and simulated uncorrelated cells, using the splatter package.

**Identification of angular progression of cycling and deuterosomal cells:** Independent sub-processing of either the cycling progenitor cells (n=759) or all the deuterosomal cells (n=2911) was done by performing batch correction and replicate merging on each cell subset, using the cell cycle-annotated genes or the combined oscillatory and transiently expressed gene sets, respectively. The circular cell trajectories in the resulting corrected PCA embedding, depicted as a black line in Fig. 3, were computed for each subset of cells using a smoothed spline passing by the center of mass of each of twelve angular bins equally distributed around the origin of the two first PCA dimensions. The angular progression of each cell along this circular cell trajectory, bound between 0 and $2\pi$, was then defined as the angle between the origin placed at (0, 1) and the cell itself, in the clockwise direction. These angular progression scores were shifted (modulo $2\pi$) in order for the 0 to align with the beginning of the cell cycle for cycling progenitors or with the beginning of the early deuterosomal cell population for deuterosomal cells. Continuous gene expression along the angular progression of either cycling progenitors (“cell cycle”) or deuterosomal cells (“MCC cell cycle variant”) was modeled using a generalized additive model using the tradeSeq package. Genes with a varying expression along either progenitor or deuterosomal cell circular trajectories were identified using the associationTest from the tradeSeq package as genes with a fold-change greater than 2 and a p-value lower than 0.01.

**RNA velocity analysis:** RNA velocity was calculated using the "dynamic" mode from scvelo software and for visualization, the RNA velocity vector field was embedded in UMAP projection of the dataset using velociraptor package. Individual RNA velocity vectors were decomposed in normal and tangential components using a coordinate plane orthogonal to each cell's angular progression.

**Gene-cyclin correlation analysis:** The pairwise Spearman correlation scores for the expression between any gene and any individual cyclin was computed. Any correlation score greater than 0.3 (with an associated p-value lower than 0.01) was then used to compute a correlation network with four nodes corresponding to each cyclin. Non-redundant sets of cyclin-coordinated genes were built by associating each gene to its most correlated cyclin.

Analysis of other MCC scRNAseq datasets

***Mouse brain during embryonic development (in vivo)***

In vivo single-cell RNA-seq data of the brain during mouse embryonic development was retrieved from La Manno et al., 2021 (https://storage.googleapis.com/linnarsson-lab-loom/dev_all.loom). Glial pre-annotated cells older than E16 were extracted and batch effects associated with "batch", "replicate", "age" and "ChipID" annotations were regressed out using regressBatches correction from batchelor package, using only marker genes identified in any of the replicates using scran package. For visualization, UMAP embedding was performed using 50 dimensions from the corrected PCA; the embedding was performed several times with a changing seed and a representative 2D projection of the dataset was used. Pre-existing cell clustering was used to re-annotate cell types and cell types irrelevant for this study (neuroblasts, oligodendrocytes, sub-commissural cells, hypendemal cells and Pdlim4+ cells) were removed. Cell re-clustering was performed on the remaining cells embedded in a SNN graph (k=5 neighbors) using the Louvain algorithm from igraph, and clusters were manually annotated using MCC differentiation markers. Putative cell cycle phases were annotated using Seurat. Genes in any given cell cluster were annotated as expressed as long as they had a log-normalized expression greater than 0.5 in more than 10% of the cells within the cell cluster.

Independent sub-processing of deuterosomal cells was done by re-embedding deuterosomal cells only in a PCA dimensional space. The same approach describe hereabove was then used to compute a circular cell trajectory and to model continuous gene expression along the angular progression of deuterosomal cells.

***Mouse tracheal embryonic cells (in vitro)***

Single-cell RNA-seq data of mouse tracheal embryonic cells in air-liquid interface (ALI) culture for 3 days was obtained from Ruiz-Garcia et al., 2019 (GEO accession ID: GSM3439924). Empty cells (detected by emtpyDrops from DropletUtils) were filtered out and only protein-coding genes were retained. The dataset was embedded in a lower PCA 50-dimension space with removed technical noise, using denoisePCA function from the scuttle package. Cell clustering was performed on cells embedded in a SNN graph using the Louvain algorithm from igraph, and clusters were manually annotated using MCC differentiation markers as well as markers for other cell types commonly found in respiratory tracts. Putative cell cycle phases were annotated using Seurat. Genes in any given cell cluster were annotated as expressed as long as they had a log-normalized expression greater than 0.5 in more than 20% of the cells within the cell cluster.

***Human airway embryonic cells (in vitro)***

Pre-processed data of single-cell RNA-seq dataset of the human airway epithelial cells in air-liquid interface (ALI) culture for 28 days (Ruiz-Garcia et al., 2019, GEO accession ID: GSM3439922) was provided by the authors. Only protein-coding genes were retained. Cell clusters, annotations and embeddings in PCA or t-SNE provided by the authors were used. Putative cell cycle phases were annotated using Seurat. Genes in any given cell cluster were annotated as expressed as long as they had a log-normalized expression greater than 0.5 in more than 10% of the cells within the cell cluster.

***Human airway epithelial cells (in vivo)***

Pre-processed data of a 500K single-cell RNA-seq dataset of human airway epithelial cells (HLCA: Human Lung Cell Atlas) (Sikkema et al. 2023) was recovered from Cellxgene (https://cellxgene.cziscience.com/collections/6f6d381a-7701-4781-935c-db10d30de293).

1,004 pre-annotated deuterosomal cells were extracted and further processed. Putative cell cycle phases were annotated using Seurat. The same approach describe hereabove was then used to compute a circular cell trajectory and to model continuous cyclin expression along the angular progression of deuterosomal cells.

***Spermatogenesis scRNAseq dataset***

Single-cell RNA-seq data of human adult spermatogenesis was obtained from Guo et al., 2018 (GEO accession ID: GSE112013). Cell annotations were retrieved from UCSC (https://cells.ucsc.edu/testis/meta.tsv) and irrelevant cells (endothelial cells, Sertoli cells, macrophages) were filtered out. Only protein-coding genes were retained. Batch effects associated with “donor” annotations were removed using fastMNN correction from batchelor package, using the 10% most variable genes. For visualization, t-SNE embedding was performed using 50 dimensions from the corrected PCA; the embedding was performed several times with a changing seed and a representative 2D projection of the dataset was used. Trajectory and pseudotime inferences were performed using PCA embedded data with slingshot, specifying start and end clusters to orientate the trajectory. Continuous gene expression along the slingshot-inferred trajectory was modeled using a generalized additive model.
